# Journey to the Center of the Mitochondria Guided by the Tail Anchor of Protein Tyrosine Phosphatase 1B

**DOI:** 10.1101/000836

**Authors:** Julia Fueller, Mikhail Egorov, Kirstin A. Walther, Ola Sabet, Jana Mallah, Markus Grabenbauer, Ali Kinkhabwala

**Author notes:** Corresponding author: Ali Kinkhabwala, Department of Systemic Cell Biology, Max Planck Institute of Molecular Physiology, Otto-Hahn-Strasse 11, 44227 Dortmund, Germany.

## Abstract

The canonical protein tyrosine phosphatase PTP1B has traditionally been considered to exclusively reside on the endoplasmic reticulum (ER). Using confocal microscopy, we show that endogenous PTP1B actually exhibits a higher local concentration at the mitochondria in all mammalian cell lines that we tested. Fluorescently labeled chimeras containing full-length PTP1B or only its 35 amino acid tail anchor localized identically, demonstrating the complete dependence of PTP1B’s subcellular partitioning on its tail anchor. Correlative light and electron microscopy using GFP-driven photo-oxidation of DAB revealed that PTP1B’s tail anchor localizes it to the mitochondrial interior and to mitochondrial-associated membrane (MAM) sites along the ER. Heterologous expression of the tail anchor of PTP1B in the yeast *S. cerevisiae* surprisingly led to its exclusive localization to the ER/vacuole with no presence at the mitochondria. Studies with various yeast mutants of conserved membrane insertion pathways revealed a role for the GET/TRC40 pathway in ER insertion, but also emphasized the likely dominant role of spontaneous insertion. Further studies of modified tail isoforms in both yeast and mammalian cells revealed a remarkable sensitivity of subcellular partitioning to slight changes in transmembrane domain (TMD) length, C-terminal charge, and hydropathy. For example, addition of a single positive charge to the tail anchor was sufficient to completely shift the tail anchor to the mitochondria in mammalian cells and to largely shift it there in yeast cells, and a point mutation that increased TMD hydropathy was sufficient to localize the tail anchor exclusively to the ER in mammalian cells. Striking differences in the subcellular partitioning of a given tail anchor isoform in mammalian versus yeast cells most likely point to fundamental differences in the lipid composition of specific organelles (e.g. affecting membrane charge or thickness) in higher versus lower eukaryotes. Fluorescence lifetime imaging microscopy (FLIM) detection of the Förster Resonance Energy Transfer (FRET)-based interaction of the catalytic domain of PTP1B with the epidermal growth factor receptor (EGFR/ErbB1) at the mitochondria revealed a strong interaction on the cytosolic face of the outer mitochondrial membrane (OMM), suggesting the presence of a significant pool of PTP1B there and a novel role for PTP1B in the regulation of mitochondrial ErbB1 activity. In summary, in addition to its well-established general localization along the ER, our results reveal that PTP1B specifically accumulates at MAM sites along the ER and localizes as well to the OMM and mitochondrial matrix. Further elucidation of PTP1B’s roles in these different locations (including the identification of its targets) will likely be critical for understanding its complex regulation of general cellular responses, cell proliferation, and diseased states.

## Introduction

The founding member of its family, protein tyrosine phosphatase 1B (PTP1B)^1,2^ (the protein product of the gene PTPN1^3^) is an important regulator of phosphotyrosine signaling in mammalian cells through its dephosphorylation of a range of substrates^4^, including the receptors for insulin, leptin, and epidermal growth factor (EGF) and their downstream substrates; the tyrosine kinases JAK2 and c-Src; and the tyrosine phosphatase SHP2. PTP1B expression has been detected in several tissues in different mammals^5^ and has been proposed as an important inhibitory target for treatment of diabetes, obesity, and cancer^6^. Its general role, particularly in cancer cell signaling, appears to be complex^7^.

PTP1B is expressed as two separate splice variants^8^, the first identified in rat brain tissue^9^ with the second later identified in human placenta^5^. In the first variant, the terminal seven amino acids of the second variant (FLFNSNT) are replaced by four amino acids (VCFH). Unlike the second variant, expression of the first is highly regulated by growth factor, with the ratiometric level of these two variants varying across different organs^8^. The subcellular localization of both variants is similar^8^. Both variants consist of an N-terminal catalytic domain and a C-terminal tail anchor^10^. A substrate “trapping mutant” of its catalytic domain^11^, the D181A mutant PTP1B^D/A^, has long provided a useful tool for understanding its catalytic mechanism as well as for enhanced detection of its interactions with substrates. PTP1B’s short (≤35 amino acid) C-terminal tail anchor was previously reported to localize it on the endoplasmic reticulum (ER)^10,12^. Exactly how the tail anchor is inserted into the membrane of the ER remains unknown, with possible contributions from the classical signal recognition particle (SRP) pathway^13^, the Sec62/63 pathway^14^, the guided entry of tail anchor proteins (GET/TRC40) pathway^15–22^, chaperone-assisted interaction^23–25^ with the ER translocon Sec61^26^, or through spontaneous insertion^27^. How these different pathways might act on PTP1B as well as how these pathways are generally tailored to engage the wide spectrum of tail-anchor-containing proteins remains largely unknown. Post-translational modification of PTP1B in an activating or inhibitory manner occurs by several mechanisms^4^, including phosphorylation (on multiple serines and tyrosines), oxidation, sumoylation, and proteolysis (calpain cleavage).

Distinct roles of different PTP1B subpopulations along the ER have been an active area of research for many years. The restriction of PTP1B to the ER has been argued as a means for regulating its interaction with plasma membrane (PM) versus endocytosed fractions of EGFR^28^. A spatial subcellular gradient of the activity of PTP1B has been proposed to account for observations of its interactions with an artificial substrate^29^. The specific roles of ER-bound PTP1B at adhesions sites^30,31^ and cell-cell junctions^32^ have also been explored. These investigations highlight important and distinct physiological roles for PTP1B subpopulations distributed across the cell.

Intriguingly, PTP1B has recently been detected within mitochondria that were extracted from rat brain tissue^33^. PTP1B’s potential presence at the mitochondria could be important for regulation of the mitochondrial phosphotyrosine proteome^34^, with important targets including several enzymes in the electron transport chain^35^, Src family kinases that localize to the mitochondria^33,36–40^, or other well-established substrates of PTP1B that also have been detected at the mitochondria like the EGF receptors ErbB1^41,42^ and ErbB2^43^ and the tyrosine phosphatase SHP2^33,40,44^. PTP1B’s mitochondrial localization, though, should be confirmed first (in particular, for more general cell lines), before further speculating on how it may reach this organelle or how its interaction there with putative as well as known substrates might affect basic mitochondrial functions.

In this manuscript, we show that PTP1B localizes to the mitochondria in multiple standard mammalian cell lines. Our detailed studies of both endogenous and overexpressed PTP1B in these cell lines establish its higher local concentration at the mitochondria than along the ER. Comparisons of full-length PTP1B with fluorescent chimeras containing only its 35 amino acid tail anchor demonstrate the complete dependence of PTP1B’s subcellular localization on its tail anchor. Higher resolution analysis using electron microscopy of DAB photo-oxidation through GFPs revealed its clear localization to the mitochondrial matrix and to mitochondria-associated membrane (MAM) sites along the ER. Heterologous expression in wild-type and mutant yeast strains allowed examination of conserved pathways that might be involved in its subcellular partitioning. These results, together with additional studies based on truncated, charge-altered, and hydropathy-altered tail anchor isoforms, paint an overall complex portrait of the dependence of PTP1B’s subcellular partitioning on the precise properties of its tail anchor. The engagement of PTP1B with its substrate ErbB1 along the ER and at the mitochondria was examined using fluorescence lifetime imaging microscopy (FLIM) of donor-labeled ErbB1 along with the acceptor-labeled trapping mutant PTP1B^D/A^ or acceptor-labeled chimeras of the catalytic domain of PTP1B^D/A^ that localized exclusively to the ER, outer mitochondrial membrane (OMM), intermembrane space (IMS), or matrix (MAT). These FLIM-based studies set the stage for future broader investigations of PTP1B’s local substrates on and within the mitochondria. PTP1B’s targeting to the mitochondria as well as its possible effects on basic mitochondrial functions will likely be essential for further elucidation of its complex roles in both normal and diseased states.

## Results

### PTP1B Localizes to the Mitochondria and Mitochondrial Interior Solely through its Tail Anchor

Recent evidence from electron microscopy and from Western blots has demonstrated the localization of PTP1B in mitochondria that were extracted from rat brain tissue^33^. To test whether this mitochondrial localization is of a more general nature, we stained multiple mammalian cell lines (COS-7, BJ Fibroblasts, HeLa, MCF7, MDCK, and HepG2) with a mitochondrial marker (MitoTracker Red CMXRos) and then fixed and permeabilized them for staining with an Alexa-488-conjugated secondary antibody that recognized a monoclonal primary antibody specific for endogenous PTP1B (Fig. 1). In these strains, endogenous PTP1B exhibited a higher local concentration at mitochondrial structures (arrows) as compared to its distribution along the ER. For the COS-7, BJ Fibroblast and HeLa cells, PTP1B accumulation at mitochondrial structures was clearly apparent in many cells, especially those having a flatter morphology with several isolated mitochondria at the cellular periphery. For MCF7, MDCK, and HepG2, it became increasingly more difficult to find mitochondria that could be cleanly separated from the more general ER staining of PTP1B. However, close examination revealed significant colocalization of PTP1B with mitochondrial structures. The challenges that we encountered in discriminating mitochondria-specific subpopulations of PTP1B from its more general ER staining in particular cell lines offers a possible explanation for the fact that its mitochondrial localization was not discovered in earlier fluorescence-based studies^10,12^.

**Figure 1.**
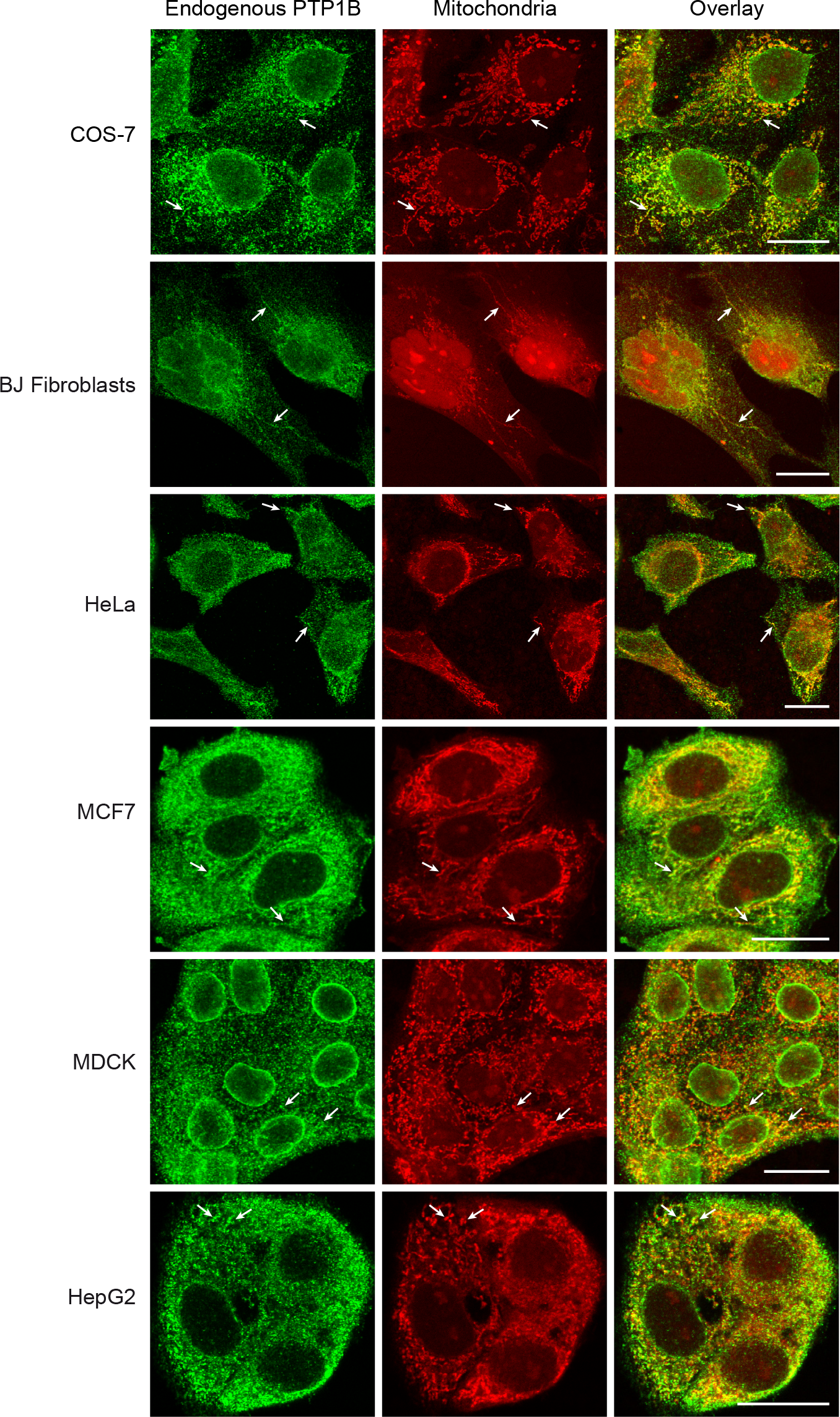
Mitochondrial localization of endogenous PTP1B in multiple mammalian cell lines. Mammalian cells were fixed with PFA and immunostained for endogenous PTP1B using an anti-PTP1B (Ab-1) mouse monoclonal primary antibody (Calbiochem) and a chicken anti-mouse secondary antibody conjugated with Alexa488 (Invitrogen). Mitochondria were stained with MitoTracker Red CMXRos (Invitrogen). COS-7, BJ Fibroblast, HeLa, MCF7, MDCK, and HepG2 cells were visualized with confocal microscopy. Representative mitochondria are marked with arrows. Mitochondrial targeting of PTP1B was easier to assess in the cells with a flatter morphology towards the top of the figure as compared with the more compact cells towards the bottom. Scale bar: 20 μm.

PTP1B’s clear accumulation in the vicinity of the mitochondria might be explained as merely the accumulation of ER-resident PTP1B at regions of the ER in close apposition to the mitochondria, so-called mitochondrial-associated membrane (MAM) sites^45–47^. To elucidate this, we prepared COS-7 cells as above but additionally expressing the ER marker mTagBFP-Sec61 (Fig. 2). A zoomed-in view of these cells shows clear accumulation of PTP1B to isolated mitochondria and mitochondrial subregions that are not in immediate proximity to the ER, proving that the observed mitochondrial localization is real and does not merely correspond to the concentration of PTP1B at a subregion of the ER.

**Figure 2.**
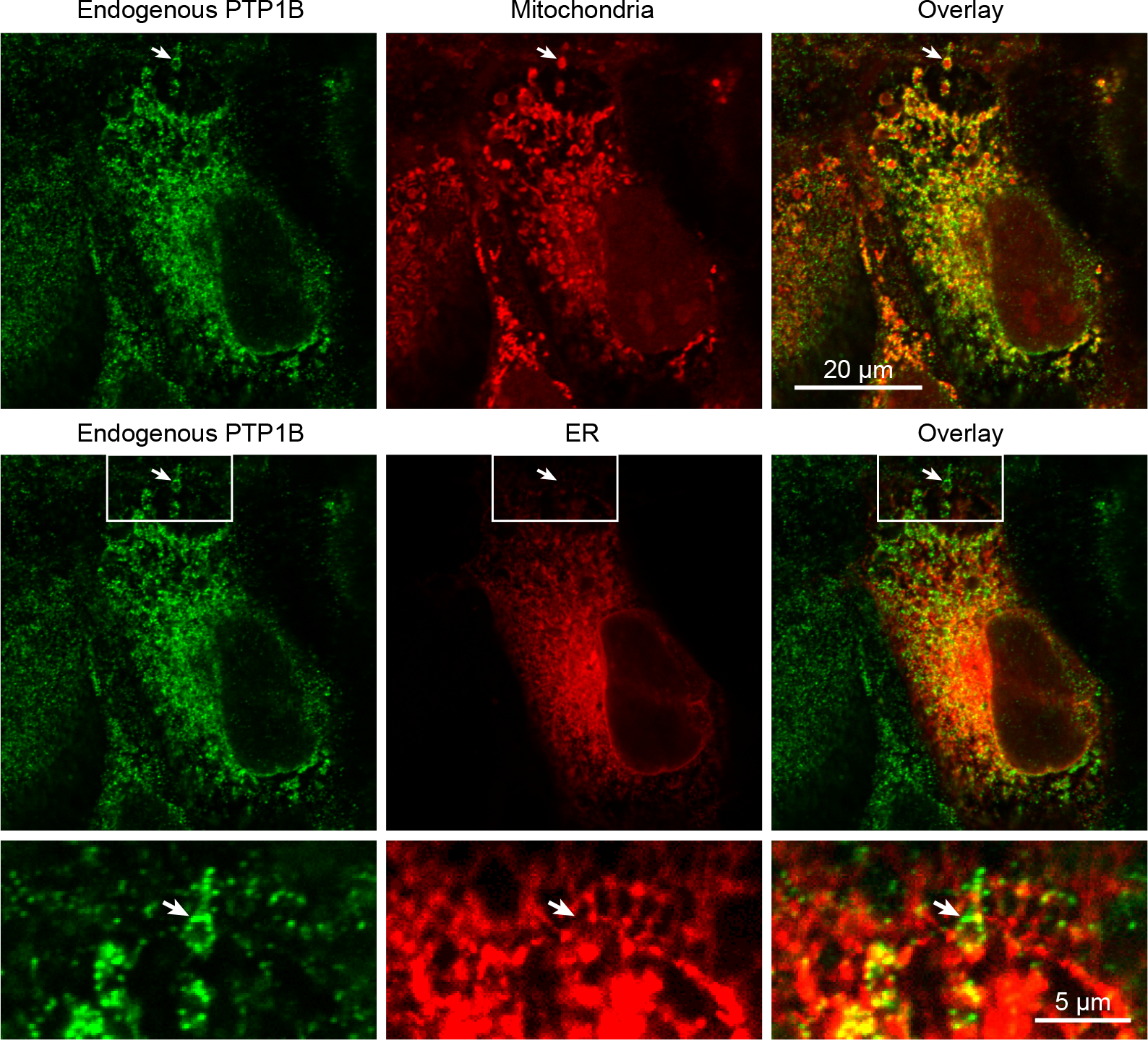
Mitochondrial versus ER localization of endogenous PTP1B in COS-7 cells. COS-7 cells were prepared for immunostaining as in Fig. 2 with the only difference being their additional expression of an ER marker (mTagBFP-Sec61). In the top row, the mitochondrial localization of endogenous PTP1B is revealed through its colocalization with the mitochondrial stain MitoTracker Red CMXRos (Invitrogen). In the second row and the third row (zoomed-in view of the white box in the second row), the discrepancy in the overlay of the PTP1B signal and the ER marker allows identification of membrane-bound PTP1B not directly associated with the ER. The structure indicated with an arrow clearly colocalizes with the mitochondria in the top row and not at all with the ER in the second and third rows. Scale bars: 20 μm and 5 μm, respectively.

Upon its expression in COS-7 cells, a chimera of mCitrine with the second splice variant of PTP1B (ending in FLFNSNT and the central focus of our current study) exhibited the same strikingly high concentration to structures coincident with the mitochondria (Fig. 3, arrow indicates individual mitochondria) as compared to its general distribution along the ER (arrowhead indicates a mitochondria-free region of the ER).

**Figure 3.**
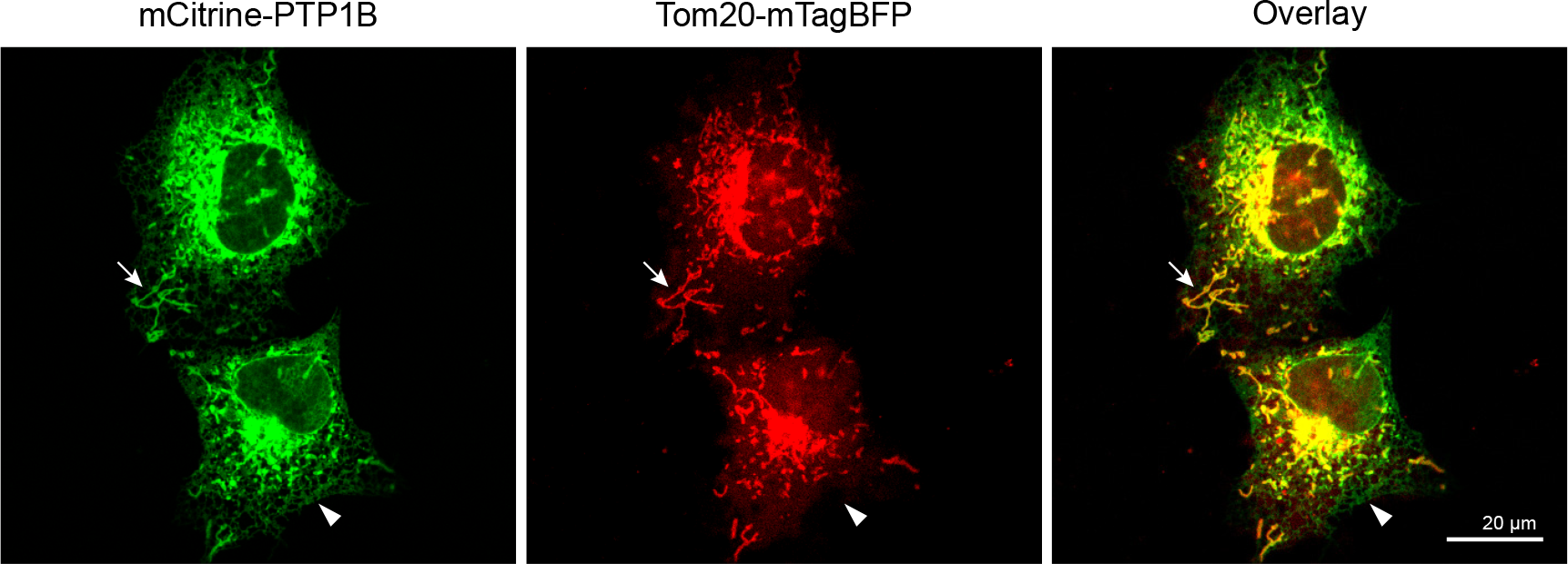
PTP1B localizes to the mitochondria in COS-7 cells. Confocal images of COS-7 cells expressing mCitrine-PTP1B along with the mitochondrial marker Tom20-mTagBFP. The mCitrine-PTP1B chimera localized to the general ER (arrowhead) and at a higher local concentration to the mitochondria (arrow). Scale bar: 20 μm.

Which domain of PTP1B is responsible for its mitochondrial partitioning? An obvious candidate is PTP1B’s ≤35 amino acid C-terminal tail anchoring domain^10,12^. Colocalization of the full-length chimera with a tail-anchor-only chimera (mCherry-PTP1Btail) yielded perfect overlap (Fig. 4A), demonstrating the complete dependence of PTP1B’s subcellular partitioning on its tail anchor. Both constructs were again heavily concentrated at the mitochondria (labeled with Tom20-mTagBFP) as compared to their distribution along the ER, which was observed for the full-length and tail-only chimeras as a faint reticular network distinct from the mitochondrial marker. We confirm as well the previous observation that the subcellular distribution of the second splice variant (PTP1Btail) is similar to that of the first variant (PTP1Btail^VCFH^), with both partitioning in the exact same manner to the mitochondria and ER (Fig. 4B).

**Figure 4.**
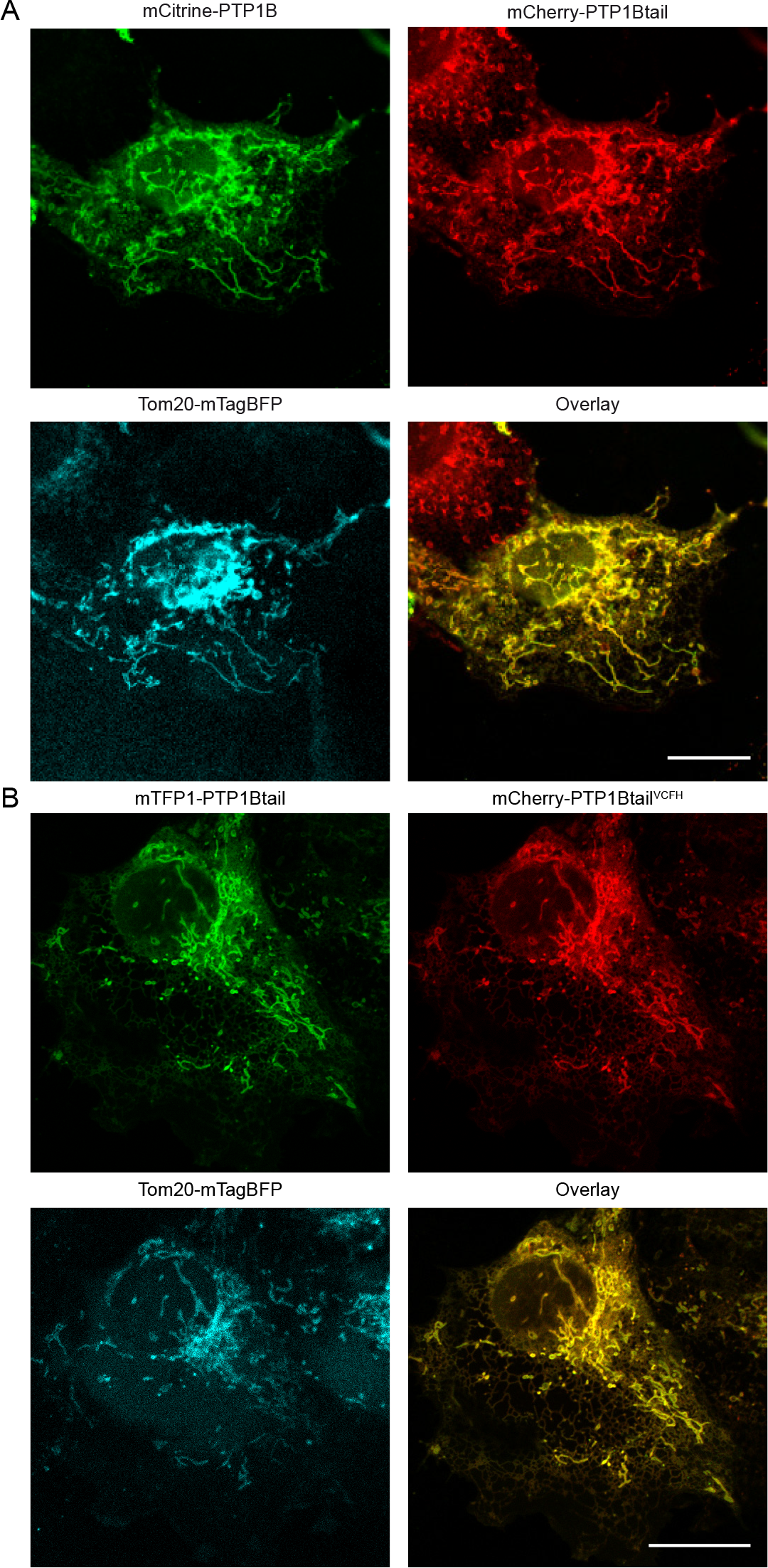
Subcellular partitioning of PTP1B is completely determined by its tail anchor. (A) COS-7 cells expressing mCitrine-PTP1B (green), mCherry-PTP1Btail (red), and Tom20-mTagBFP (blue) were visualized by confocal microscopy. The lower right image represents the overlay of the mCitrine-PTP1B (green) and mCherry-PTP1Btail (red) images. (B) COS-7 cells expressing mTFP1-PTP1Btail (green), mCherry-PTP1Btail^VCFH^ (red), and Tom20-mTagBFP (blue) were visualized by confocal microscopy. The lower right image represents the overlay of the mTFP1-PTP1Btail (green) and mCherry-PTP1Btail (red) images. Scale bar: 20 μm.

Further dynamic verification of the colocalization of full-length PTP1B with its tail anchor is shown in Movie 1 for live COS-7 cells. Pulsed-interleaved excitation (PIE, see Materials and Methods) was used to acquire essentially simultaneous (down to 25 ns) dual-color images of COS-7 cells expressing mTurquoise-PTP1B along with mCherry-PTP1Btail. Both constructs exhibited perfect, and therefore unchanging, overlap over the entire duration of the movie (6 minutes).

To examine the precise submitochondrial localization of PTP1B, we used electron microscopic analysis of GFP-driven DAB photo-oxidation (Fig. 5, see Materials and Methods). Correlative microscopy of a full-length chimera (mTurquoise-PTP1B) or a tail-anchor-only chimera (mTFP1-PTP1Btail) revealed their localization to the mitochondrial matrix and notable absent from the intracristal spaces (and likely the entire soluble part of the intermembrane space, IMS) in COS-7 cells. Its localization to the OMM or IMM was unclear due to the already significant general staining of the lipids in these membranes (see control cells, Figs. 5C and 5F). PTP1B’s distributed localization throughout the matrix suggests a significant soluble fraction in this region. Our results generally confirm prior studies based on immunogold staining of mitochondria extracted from rat brain cells, in which endogenous PTP1B was also localized to the mitochondrial interior^33^. The higher labeling efficiency possible with photo-oxidation (as compared with immune-electron microscopy) allowed visualization of its distributed localization throughout the mitochondrial matrix and its complete absence in the intracristal spaces of the IMS. Electron microscopy of the tail-anchor-only chimera (Fig. 5E) importantly demonstrated the complete dependence of PTP1B’s submitochondrial localization on its tail anchor. Intriguingly, our photo-oxidation-based images also revealed strong staining at putative MAM sites along the ER for both the full-length and tail-anchor-only chimeras (white arrows in Figs. 5B and 5E). Such accumulations point to a possible mitochondrial-targeting mechanism consisting of direct transfer from the ER to the mitochondria at these sites. Further trafficking through the OMM/IMM to the mitochondrial matrix could take place through interaction with the outer and inner membrane translocase complexes, either completely unassisted or through the mediation of specific chaperones. Previous immunogold-stained EM images of other phosphotyrosine regulators (Lyn^36^, c-Src^37^, ErbB1^42^, and ErbB2^43^) have indicated their presence as well *within* the mitochondria. How all of these important regulators of phosphotyrosine signaling manage to pass through the OMM (and IMM?) to the mitochondrial interior remains unclear.

**Figure 5.**
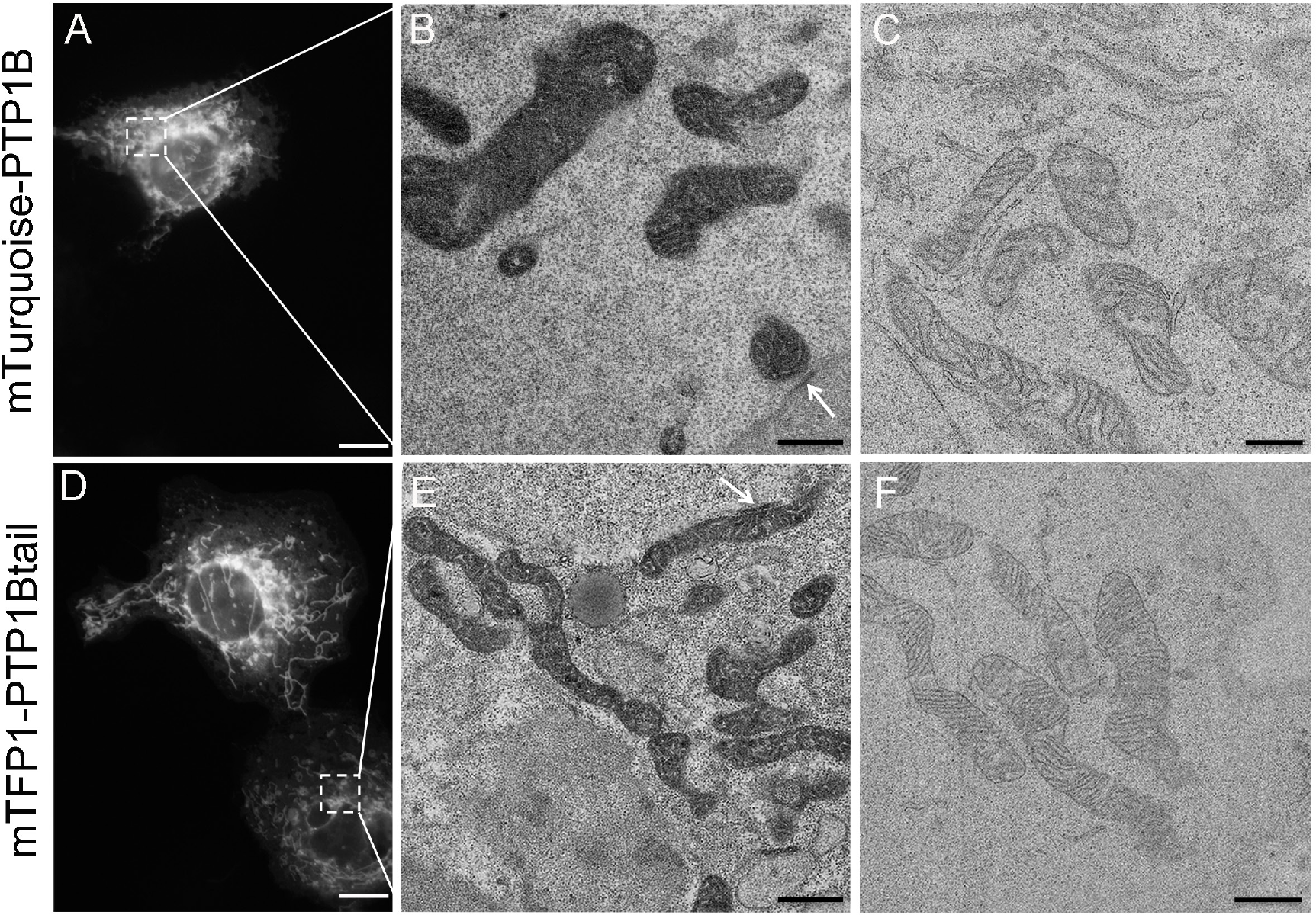
Correlative microscopy using photo-oxidation of DAB through fluorescently labeled PTP1B or its tail anchor expressed in COS-7 cells. (A) Widefield fluorescence image of a COS-7 cell expressing mTurquoise-PTP1B. (B) Correlated electron micrograph of the region indicated in A after photo-oxidation. Specific DAB staining was observed largely throughout the matrix and was notably absent from the intracristal spaces. The arrow indicates a region of higher staining along the nuclear ER that is in direct apposition to a mitochondrion and therefore consistent with a MAM site. (C) Electron micrograph of mitochondria in a nearby untransfected cell as a negative control. (D) Widefield fluorescence image of COS-7 cells expressing mTFP1-PTP1Btail. (E) Correlated electron micrograph of the region indicated in D after photo-oxidation. As for B, specific DAB staining was observed throughout the matrix and was absent within the intracristal spaces. The arrow indicates an extended region of higher staining in direct apposition to a mitochondrion and therefore consistent with a MAM site on the general ER. (F) Electron micrograph of mitochondria in a nearby untransfected cell as a negative control. Scale bars: 10 μm (A,D), 500 nm (B,C), or 1 μm (E,F).

### Investigation of Tail Anchor Targeting Pathways that May Regulate the Subcellular Partitioning of PTP1B

Our above results collectively reveal the tail-anchor-dependent subcellular partitioning of PTP1B to the ER (including its accumulation at MAM sites) and the mitochondria (including the mitochondrial interior). Tail anchor (TA)-containing proteins, which by definition contain a C-terminal hydrophobic TMD (dark blue in the cartoon and upper box in Fig. 6), can be targeted to the ER and the mitochondria via multiple pathways as depicted in Fig. 6, with the emerging view that the overall hydropathy of the TMD plays an important role in determining pathway specificity^48^. From high to low overall hydropathy, the tail anchor can be targeted to the ER by the classical SRP pathway^13^, the Sec62/63 pathway^14^, the GET/TRC40 pathway^15–22^, more general chaperones^23–25^, or spontaneous insertion^27^. For all of these pathways, the Sec61 translocon likely plays either an essential or important role (even for spontaneous insertion). Assisted (via chaperones^49^) or unassisted insertion into the OMM or into the IMS most likely involves the translocase of the outer membrane (TOM complex), with the translocase of the inner membrane (TIM complex) similarly facilitating insertion into the IMM or into the mitochondrial matrix^50,51^. Spontaneous insertion into the OMM has also been reported for several tail anchor proteins^52,53^. A more complex route could be through transfer from the ER to the mitochondria at MAM sites along the ER^45^ (already discussed above with respect to our observed accumulation of PTP1B at MAM sites, Fig. 5), which are anyway sites of dynamic lipid transfer between these organelles^47^.

**Figure 6.**
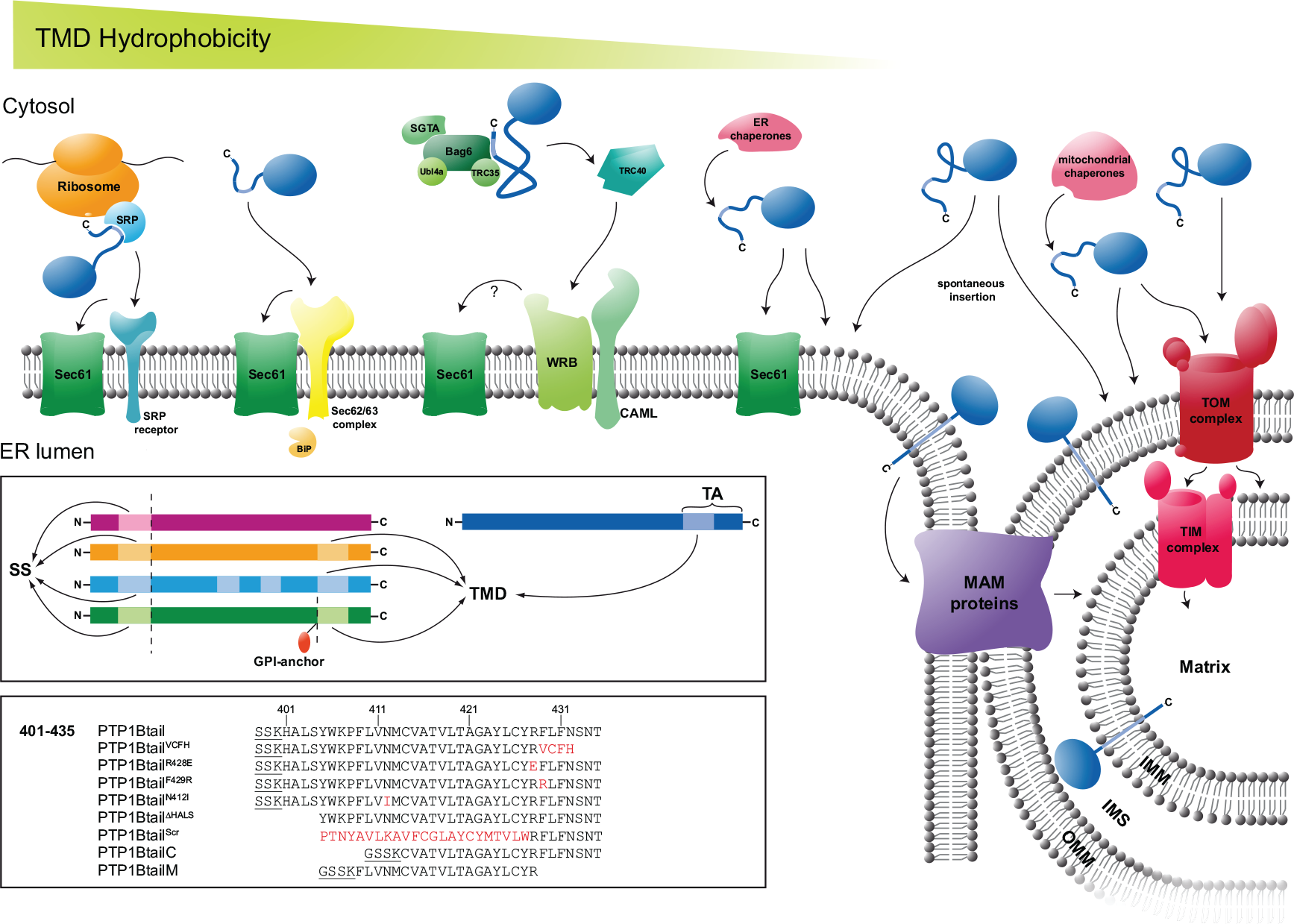
Possible pathways contributing to the subcellular targeting of the tail anchor protein PTP1B. See text for further description.

Unlike tail anchor proteins, other proteins rely on an additional (typically cleavable) N-terminal signal sequence^54^ (SS) for successful insertion into the ER membrane or for passage into the lumen. These SS-containing proteins can be divided into the following classes also displayed in the upper box in Fig. 6: magenta (proteins containing *only* the N-terminal SS), orange (proteins additionally containing a single putative C-terminal TMD), light blue (proteins additionally containing multiple TMDs), and dark green (proteins additionally containing a single putative C-terminal TMD that is cleaved in the ER lumen, with a GPI anchor attached to the new C terminus).

The lower box in Fig. 6 lists the sequences of the wild-type 35 amino acid tail (PTP1Btail) and multiple isoforms studied in this manuscript: charge-altered isoforms (PTP1Btail^R428E^, PTP1Btail^F429R^), hydropathy-altered isoform (PTP1Btail^N412I^), N-terminally truncated (PTP1Btail^ΔHALS^) and scrambled (PTP1Btail^Scr^) isoforms, and further truncated isoforms (PTP1BtailC and PTP1BtailM). Residues that link these tail isoforms to the N-terminal fluorophore sequence are underlined (when present).

To isolate which pathway(s) might be at work in targeting PTP1B to the ER and to the mitochondria, we employed the genetically pliable yeast *S. cerevisiae* to study these largely conserved tail-anchor targeting pathways. In yeast, a “hydropathic code” appears to be at work^55^, with clear differences in overall hydrophobicity of the tail anchors of proteins targeted to the various organelles (Fig. 7). The Kyte-Doolittle hydropathies^56^ of a relatively complete set of the tail anchor-containing proteins in yeast^55^, along with the class of GPI-anchored proteins like Gas1^57^ that contain a putative TMD at their C-terminus before cleavage and attachment of the GPI anchor (Fig. 6), are displayed in Fig. 7. Organellar localizations are based on observations of the full-length proteins. The pattern already revealed by this representation is striking, with the clear difference in hydropathy of the mitochondrial versus ER proteins of particular significance for the present study. The tail anchor of PTP1B is shown in black in all of these figures. Based on hydropathy alone, therefore, we predicted that the tail anchor would preferentially localize to the mitochondria or to the nuclear envelope.

**Figure 7.**
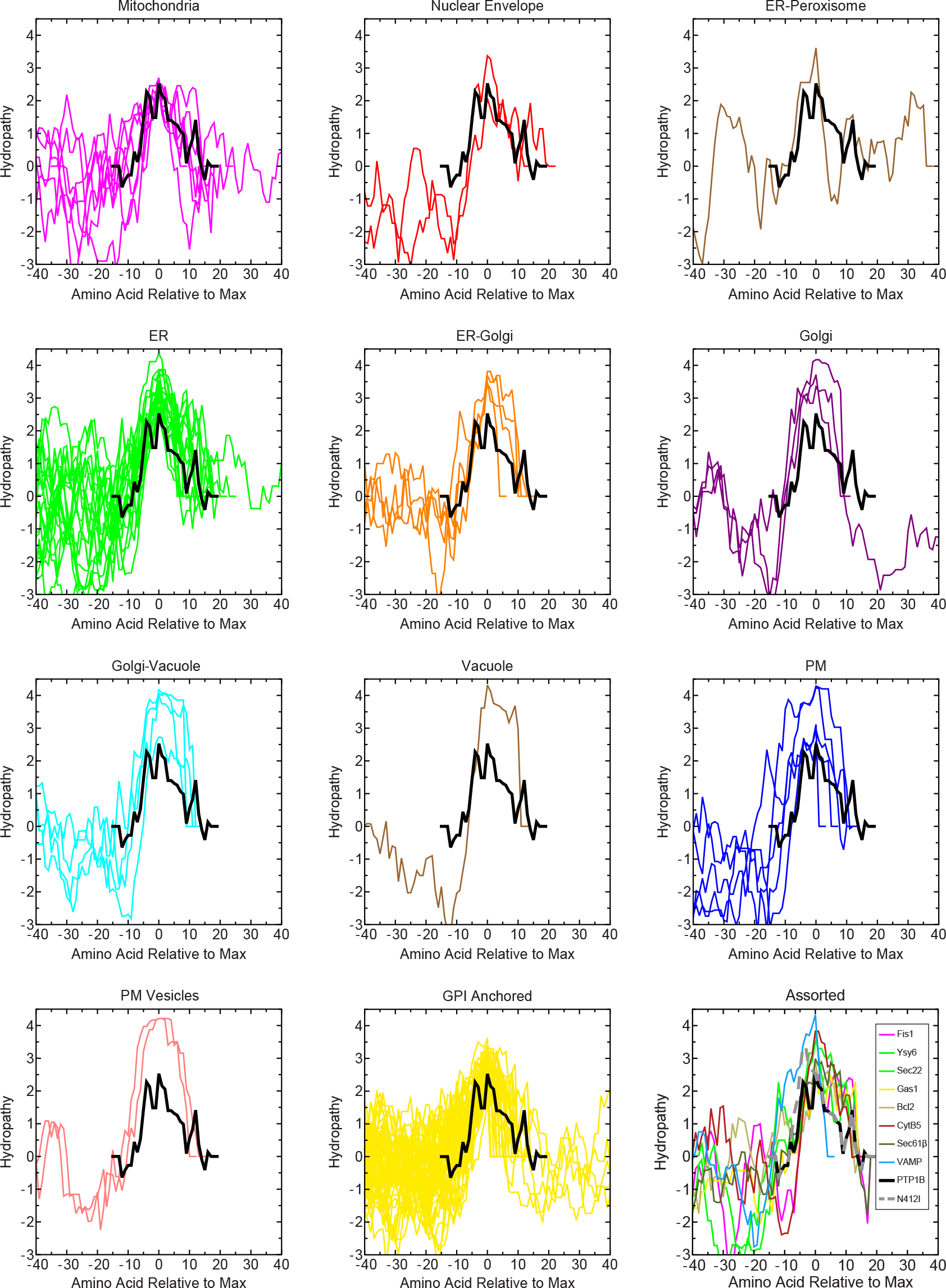
Predicted subcellular targeting of the PTP1B tail anchor in yeast based on its hydropathy. A library of previously identified C-terminal tail-anchor-containing proteins in yeast is displayed based on the list of Burri & Lithgow^55^ (with additional inclusion of Gem1^96^ in the mitochondria panel) along with the list of 56 GPI-anchored proteins given in Ast et al.^57^, which were removed from the list of Burri & Lithgow due to their GPI anchorage. The hydropathies of the C termini of these proteins are displayed based on the Kyte-Doolittle method^56^ (boxcar smoothing of n=7), with all protein sequences centered at the amino acid position corresponding to peak hydropathy. The hydropathy profile of the PTP1B tail anchor is shown in all figures (black). In the bottom right panel, the hydropathy profiles of canonical tail anchor proteins from yeast (Fis1, Ysy6, Sec22, Gas1) and from mammalian cells (Bcl2, CytB5, Sec61β, VAMP-1B) are shown, as well as the high hydropathy tail isoform PTP1B^N412I^ (N412I, which has not been shifted to its maximum hydropathy, but is instead in register with the PTP1B tail).

To our surprise, however, a tail-anchor-containing chimera (yemCitrine-PTP1Btail) localized to the ER and the vacuolar membranes upon its heterologous expression in yeast (Fig. 8), with no significant concentration at the mitochondria or any other organelles.

**Figure 8.**
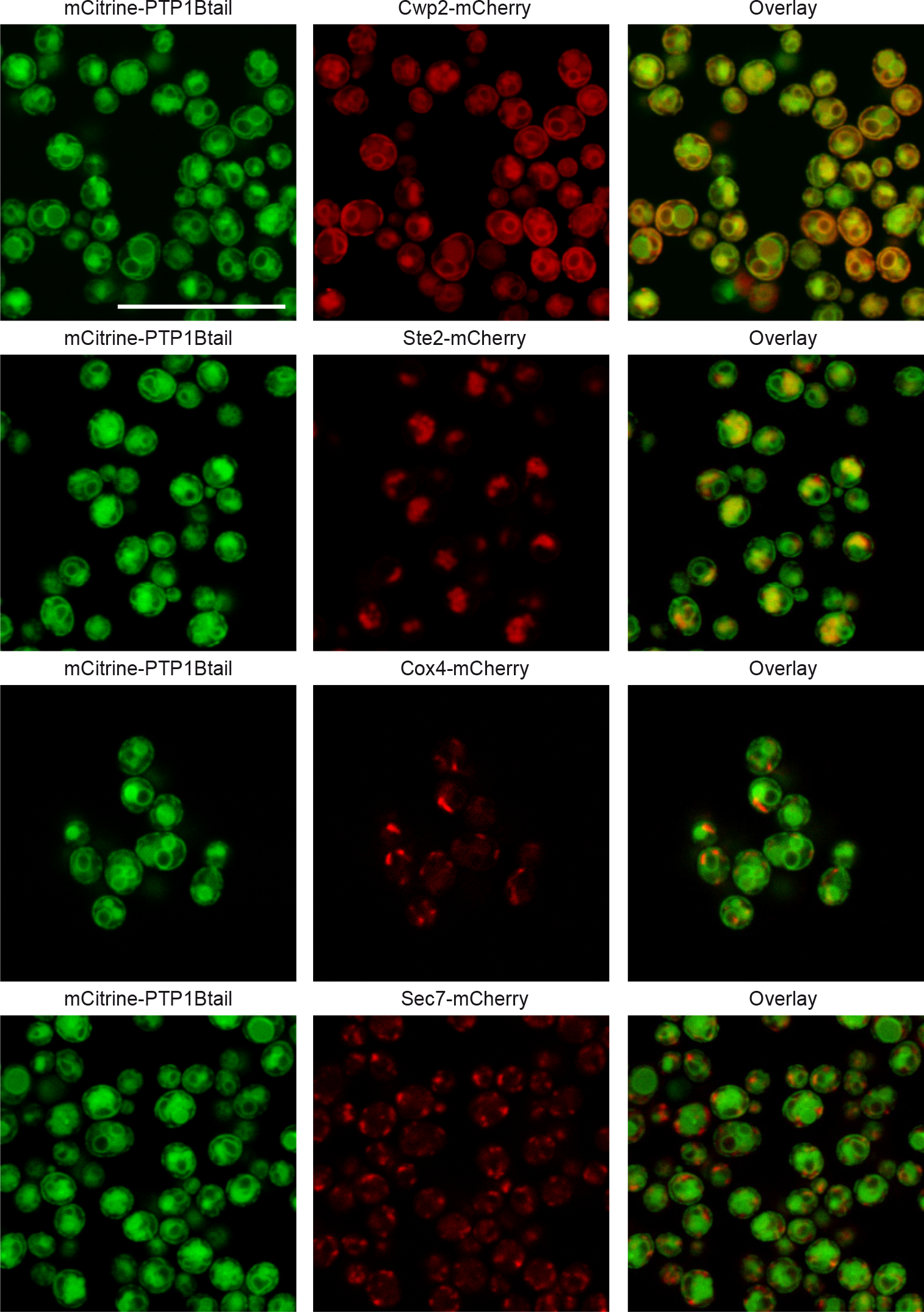
Subcellular partitioning of the tail anchor of PTP1B in the yeast *S. cerevisiae* (ESM356-1 background strain^91^). Confocal microscopy of specific strains that chromosomally expressing yemCitrine-PTP1Btail and markers for either the ER (Cwp2-mCherry), vacuole (Ste2-mCherry), mitochondria (Cox4-mCherry), or Golgi (Sec7-mCherry). Scale bar: 20 μm.

What could account for this discrepancy? One of the principal differences between yeast membranes and higher eukaryotic membranes is the production and incorporation of ergosterol versus cholesterol. Membranes that incorporate ergosterol are more ordered and therefore thicker and less fluid than those that incorporate the same concentration of cholesterol^58^ and could therefore account for this difference in stability of the tail anchor’s organelle-specific insertion in yeast versus mammalian cells. To test this, we used a recently reported yeast strain in which intracellular ergosterol is replaced by cholesterol through exchange of the two terminal enzymes in the ergosterol production pathway with their cholesterol-specific counterparts^59^. Localization of the tail anchor of PTP1B in this mutant strain (cholesterol) was identical to that in its wild-type (ergosterol) counterpart (Fig. 9), indicating no influence on the presence of specific sterols on the subcellular targeting of the tail anchor of PTP1B.

**Figure 9.**
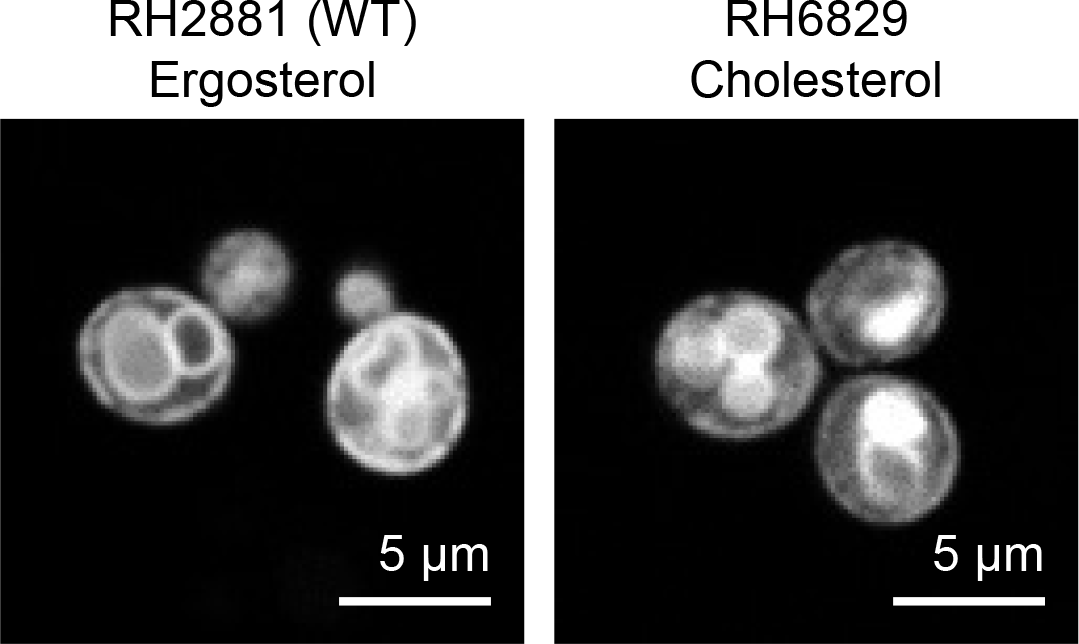
Dependence of the targeting of the tail anchor of PTP1B on the exact sterol. A wild-type strain that produces ergosterol (RH2881) as well as a mutant strain that produces cholesterol as its dominant sterol (RH6829), with membrane composition therefore more similar to mammalian cells, were transformed with the plasmid p415-yemCitrine-PTP1Btail. In both strains, yemCitrine-PTP1Btail localized to the perinuclear/cortical membranes of the ER and to the vacuolar membrane. Scale bars: 5 μm.

To investigate the specific mechanism(s) that underlie the membrane targeting of the tail anchor in yeast, we examined mutant yeast strains that were deleted in important proteins involved in the various pathways for tail anchor targeting (Fig. 10).

**Figure 10.**
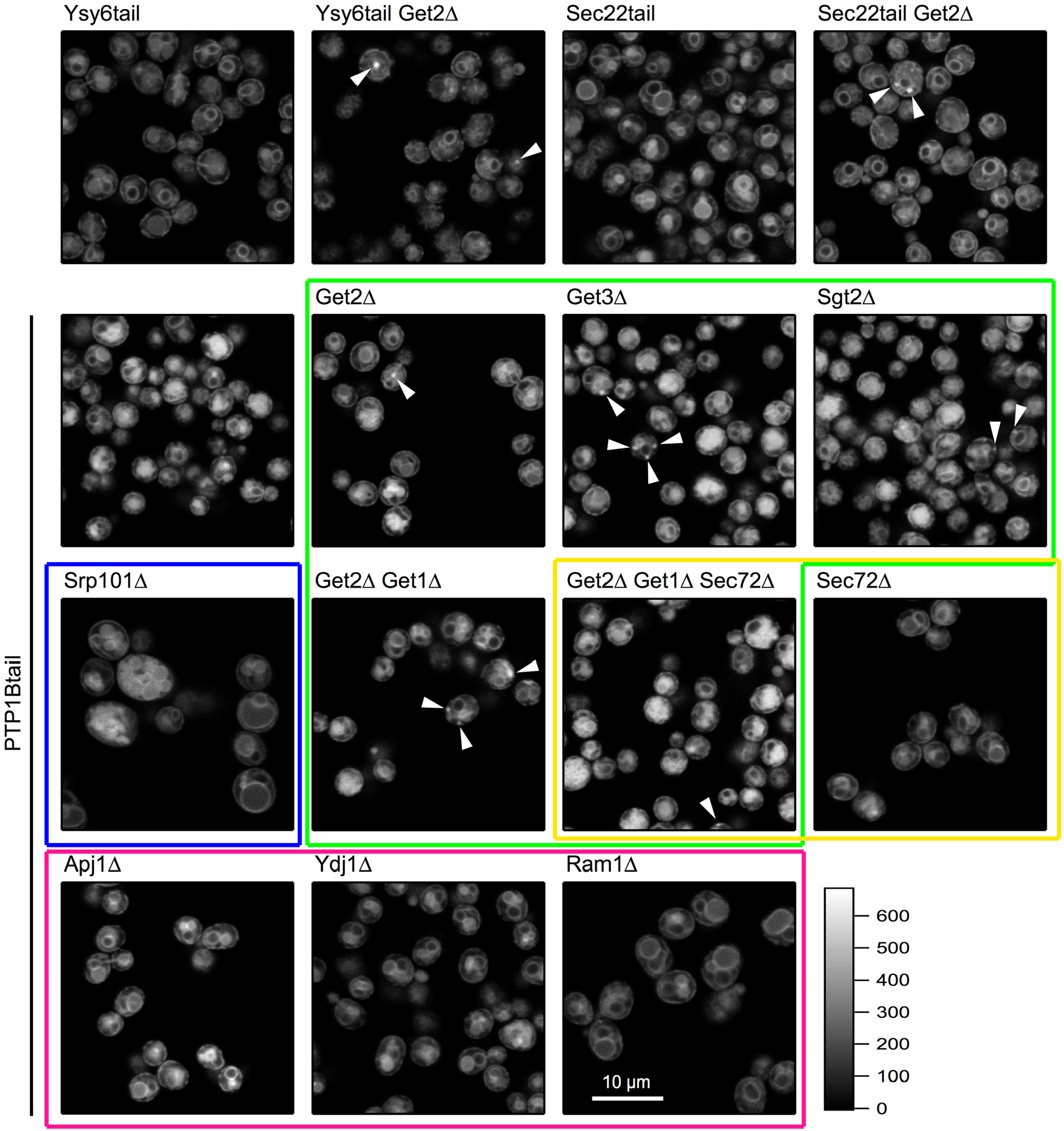
Systematic study of the potential pathways responsible for the subcellular targeting of the tail anchor of PTP1B in yeast. We used an aggregate-formation assay to examine the impact of deletion of the various insertion pathway components in yeast. Aggregate formation in strains expressing tail anchors from Ysy6 and Sec22 upon deletion of the GET pathway component Get2 was used as a control. Comparing wild-type with Get2Δ deletion strains that expressed yemCitrine-Ysy6tail or yemCitrine-Sec22tail, we observed visible aggregates in a fraction of the Get2Δ cells that were consistent with previous findings (though the bulk of these proteins still managed to insert properly). We observed similar aggregates in cells expressing yemCitrine-PTP1Btail. Slightly more aggregates were observed in Get3Δ cells and significantly fewer in the Sgt2Δ cells of this strain. A strain in which both Get2 and Get1 were deleted was similar to the Get2Δ deletion strain. The green box indicates all strains in which one or more GET pathway components were deleted. Deletion of the α subunit of the SRP receptor (Srp101) led to a much larger cell phenotype (blue box), but no change in the ER and vacuolar partitioning of the PTP1B tail anchor. Deletion of Sec72 (yellow box), an essential factor of the Sec62/63 pathway, also did not alter the localization of the PTP1B tail anchor. Deletion of Sec72 in the Get2Δ/Get1Δ strain did not further alter the localization of the PTP1B tail anchor beyond that observed in the Get2Δ/Get1Δ strain. Finally, deletion of the chaperones Apj1 and Ydj1 (as well as the farnesyl transferase of Ydj1, Ram1) did not alter PTP1B’s subcellular localization (red box). The intensity scale shown at the bottom right and indicating photon counts/pixel is applicable to all images, as is the scale bar shown in the bottom right panel. Scale bar: 10 μm.

A major mechanism for tail insertion is through the GET/TRC40 pathway^15–19,21,22^. To examine its role in the insertion of the tail anchor of PTP1B, we systematically deleted proteins involved in this pathway in a yeast strain chromosomally expressing yemCitrine-PTP1Btail. Deletions of ER-resident Get2 (functional mammalian homologue CAML^60^), cytosolic chaperone Get3 (mammalian homologue TRC40^16^), the chaperone-interacting protein Sgt2 (mammalian homologue SGTA^61^), and/or the ER-resident Get1 (mammalian homologue WRB^62^) led to the formation of cytosolic aggregates, indicating the involvement of the GET pathway. Similar aggregates were observed under the same experimental conditions for the tail anchors of Ysy6 and Sec22, which were previously identified as GET pathway targets using the same phenotypic assay^17,19^. Note that there is no increase in the level of cytosolic aggregates in the Get2Δ/Get1Δ strain over that observed in the Get2Δ strain, in line with the observation that Get1 is unable to insert in the ER in a Get2Δ background^63^. We caution that, despite these clear phenotypes, the GET pathway may still not be *directly* responsible for insertion of any of these tail anchors. Aggregate formation in these mutants could be nucleated by a distinct set of proteins dependent on the GET pathway; once such aggregates are formed, they could recruit other tail anchor proteins (including possibly PTP1B, Ysy6, or Sec22) that are not directly dependent on the GET pathway for their insertion (e.g. normally spontaneously inserted). In all of these GET pathway mutants, a significant fraction of the tail anchors of PTP1B, Ysy6, and Sec22 indeed still manages to reach the ER, suggesting spontaneous insertion as the likely dominant mechanism.

Evidence for a role of the SRP pathway in the post-translational targeting to the ER of some tail anchor proteins like the β subunit of Sec61 and synaptobrevin has been previously reported^64^. To abrogate the SRP pathway, we deleted SRP101, the essential α subunit of the SRP receptor. Deletion of SRP101 results in a viable strain^65^ but one with six-fold slower growth and a roughly three-fold increase in cell size (Fig. 10). In such a mutant, the tail anchor still localized entirely to endomembranes consistent with the ER and vacuole.

To investigate the contribution of the Sec62/63 pathway, we deleted the essential component Sec72 of this pathway in yeast. The tail anchor of PTP1B inserted normally in this strain, with the phenotype of a strain triply deleted for Get2/Get1/Sec72 identical to that of the double deletion Get2/Get1 (Fig. 10). These results were in line with similar observations of heterologously expressed mammalian cytochrome B5 (CytB5) in yeast, which also showed no dependence on the Sec62/63 pathway for its insertion (through use of temperature-sensitive mutants of the Sec62/63 pathway^66^).

A possible role for general chaperones in the insertion of the similarly tail anchored CytB5 has previously been hypothesized based on oxidation studies^67^. Deletions of the chaperones Apj1 (DnaJ-family) and Ydj1 (Hsp40), as well as the farnesyl transferase Ram1 of Ydj1 were previously reported to affect insertion of GPI-family mutants through the observation of visible aggregates in many cells^57^. These GPI-family proteins contain C-terminal regions with similar hydropathy to the PTP1B tail anchor (Fig. 7), making their particular chaperones interesting targets for assessing their effect on PTP1B’s membrane insertion. Indeed, a possible role of Hsp40/Hsp70 chaperone-assisted insertion of PTP1B into the ER of mammalian cells has previously been reported^25^. For the PTP1B tail, however, deletion of Apj1, Ydj1 (Hsp40), and Ram1 did not at all affect its localization to the ER and vacuolar membranes.

To conclude, the tail anchor of PTP1B is possibly inserted with the help of the GET pathway or other as yet unidentified factors (e.g. chaperones), but spontaneous insertion (as previously shown in vitro^27^) appears to be the most likely mechanism.

### Dependence of the Subcellular Targeting of PTP1B on the Exact Composition of its Tail Anchor

The length of the hydrophobic domain of a tail anchor protein can determine its membrane specificity^68,69^. To explore the role of tail anchor length on the subcellular targeting of PTP1B, we examined truncations of the tail from either end. A truncation of the N-terminal sequence (HALS) exhibited an identical partitioning to the mitochondria/ER as for the full-length tail (Fig. 6, see Fig. 15A below), proving that the 31 amino acids at the C-terminus of PTP1B are already sufficient to account for its localization.

Further truncation of the N-terminus, resulting in the PTP1BtailC construct of only 22 amino acids (Fig. 6), was previously shown to be sufficient for its ER targeting^12^. We found, however, that the PTP1BtailC construct localized it not only to the ER, but also to the Golgi and cytosol in COS-7 cells, with no presence at the mitochondria (Fig. 11A,B). The PTP1BtailC chimera was also found at rapidly-moving vesicles (punctate structures in Figs. 11A and 11B). Unexpectedly, in yeast the PTP1BtailC chimera localized to the ER *and* the mitochondria, with no localization at the Golgi (Figs. 11C–E).

**Figure 11.**
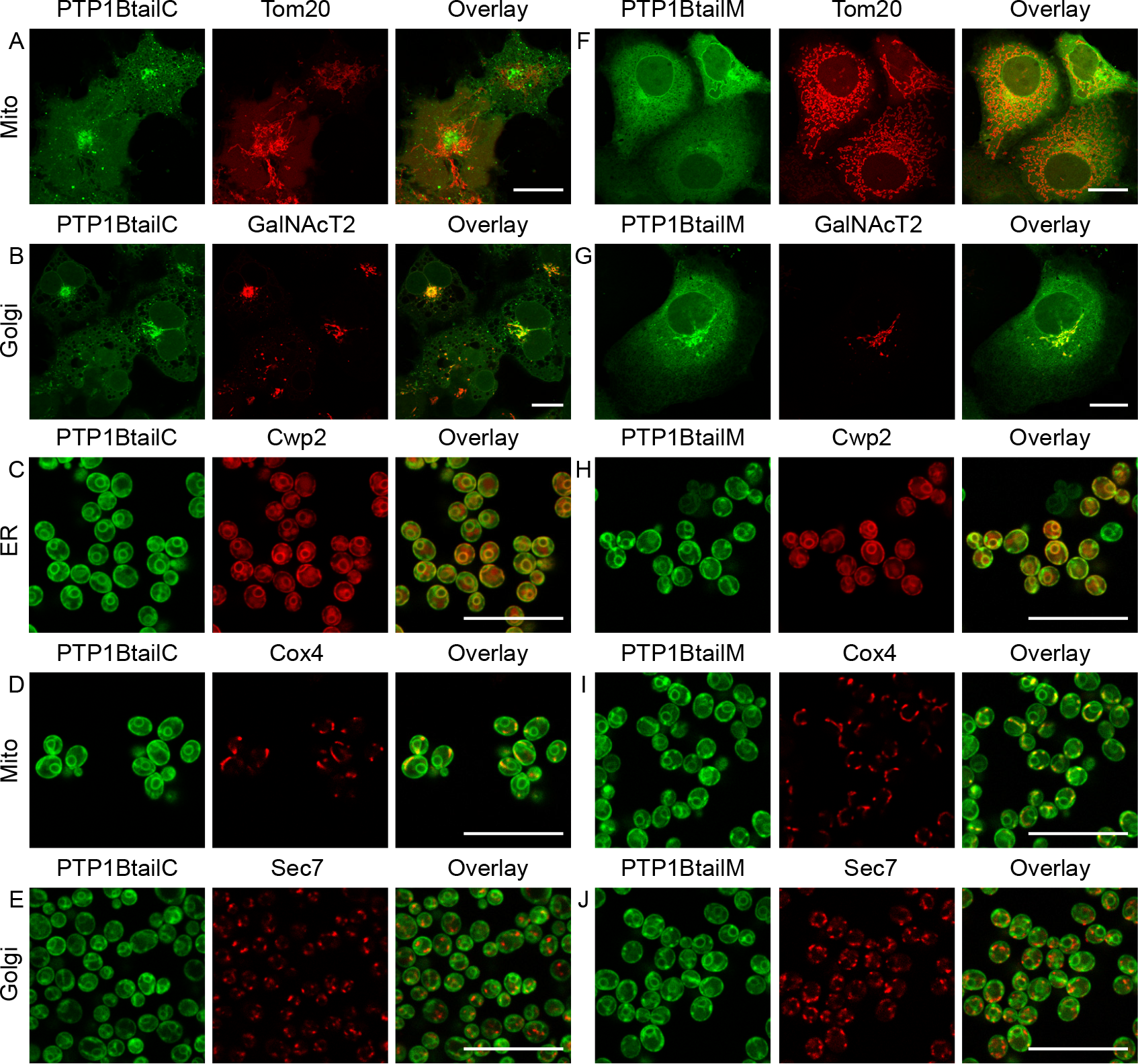
Localization of N- and/or C-terminal truncations of the tail anchor of PTP1B in COS-7 cells and in yeast. (A–E) Coexpression of the N-terminally truncated and fluorophore-labeled PTP1BtailC (Fig. 6) in COS-7 cells (mCherry-PTP1BtailC) along with either the mitochondrial marker Tom20-mTagBFP (A) or the Golgi marker GalNAcT2-mTurquoise (B); and in yeast cells (yemCitrine-PTP1BtailC) along with either the ER marker Cwp2-mCherry (C), the mitochondrial marker Cox4-mCherry (D), or the Golgi marker Sec7-mCherry (E). (F–J) Similar coexpression of the N- and C-terminally truncated and fluorophore-labeled PTP1BtailM (Fig. 6) in COS-7 cells (mCherry-PTP1BtailM, F and G) and in yeast (yemCitrine-PTP1BtailM, H–J). Scale bars: 20 μm.

Another tail anchor mutant spanning the “middle” putative-TMD-containing region of the wild-type tail anchor (PTP1BtailM, Fig. 6) was also previously shown to be sufficient for its ER targeting^12^. As for the PTP1BtailC anchor, we found that this tail anchor localized not only to the ER, but also to the Golgi and cytosol in COS-7 cells, with again no presence at the mitochondria (Figs. 11F and 11G). Unlike the PTP1BtailC isoform, though, no further localization to rapidly moving vesicles was observed. In yeast, the PTP1BtailM chimera localized identically to the PTP1BtailC isoform to the ER and the mitochondria but not the Golgi (Figs. 11H–J).

In COS-7 cells and in yeast cells, both the PTP1BtailC and PTP1BtailM isoforms were equally able to reach the ER; however, the additional difference in localization to the Golgi (COS-7) or to the mitochondria (yeast) is puzzling. In the case of spontaneous insertion, this could be explained by differences in lipid composition of both the Golgi and the mitochondria in COS-7 versus yeast. Such organellar differences in lipid composition could affect the fluidity of the membrane as well as the width of the lipid bilayer^70^ (determined by the properties of the acyl chains, in particular their lengths), both of which could alter the retention of these tail anchor isoforms.

To summarize, truncations of the 35 amino acid tail anchor from its N- and/or C-termini, while preserving the ability to insert into endomembranes^12^, can nevertheless generate dramatic differences in *specific* subcellular partitioning that intriguingly differ for yeast versus higher eukaryotes.

The C-terminal charge of tail anchors has been shown in the past to affect the targeting of tail anchor proteins. For PTP1B, the arginine at position 428 out of 435 contributes a single positive charge at the C terminus. To examine the effect of the presence of this charge on the targeting of PTP1B, we expressed mutants for which this positive charge is replaced with a negative charge (PTP1Btail^R428E^, Fig. 6) or augmented by an additional neighboring positive charge (PTP1Btail^F429R^, Fig. 6). Conversion to a negative charge led to complete restriction of the tail anchor to the ER with no detectable mitochondrial presence in COS-7 cells (Fig. 12A); in yeast, the negative tail anchor localized largely to the ER and only somewhat to the vacuole (Fig. 12B) in contrast to the more equal ER/vacuolar distribution of the wild-type tail (Fig. 8). Augmentation of the C-terminal positive charge led to complete targeting of the tail anchor to the mitochondria in COS-7 cells (Fig. 12C); in yeast, this tail anchor was largely redirected to the mitochondria but also still somewhat present on the ER/vacuole (Fig. 12D).

**Figure 12.**
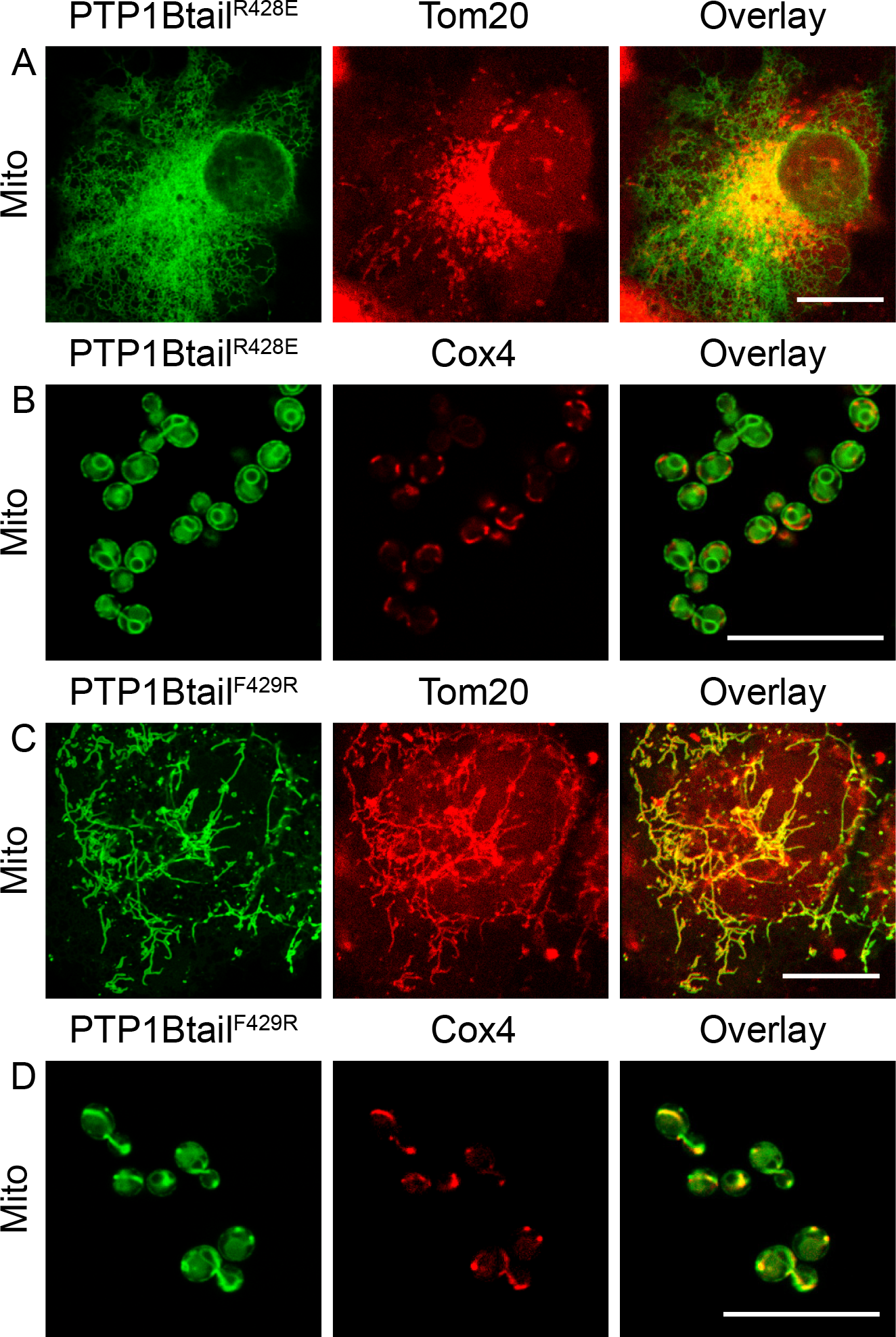
Localization of charge-altered isoforms of the tail anchor of PTP1B in COS-7 cells and in yeast. (A,B) Coexpression of the fluorphore-labeled, negatively charged tail isoform PTP1Btail^R428E^ (Fig. 6) in COS-7 cells (mCherry-PTP1Btail^R428E^) along with the mitochondrial marker Tom20-mTagBFP (A) and in yeast cells (yemCitrine-PTP1Btail^R428E^) along with the mitochondrial marker Cox4-mCherry (B). (C,D) Coexpression of the fluorophore-labeled, highly positively charged tail isoform PTP1Btail^F429R^ (Fig. 6) in COS-7 cells (mCherry-PTP1Btail^F429R^) along with the mitochondrial marker Tom20-mTagBFP (C) and in yeast cells (yemCitrine-PTP1Btail^F429R^) along with the mitochondrial marker Cox4-mCherry (D). Scale bars: 20 μm.

These results are consistent with previous results on the tail anchor of CytB5. For CytB5, the wild-type tail has a net charge of −1, which localizes it exclusively to the ER^71^. A CytB5 isoform that is truncated at its C terminus, generating a +1 charged tail, localizes as well to the mitochondria^72^. Exclusive mitochondrial localization is accomplished through addition of another positive charge to this isoform, generating a +2 net charge^73^. Other tail anchor proteins for which C-terminal positive charge plays a significant role in their mitochondrial targeting include synaptobrevin/VAMP-1B^74^ and OMP25^69^ in mammalian cells and Fis1^53^ in yeast.

Based on the yeast “hydropathic code” (Fig. 7), which shows a clear preference of tail anchors with low hydropathy for the mitochondria and with high hydropathy for the ER, we hypothesized that an increase in the hydropathy of the wild-type tail anchor in mammalian cells might be sufficient to shift it away from the mitochondria to the ER. Indeed, a tail anchor with significantly higher hydropathy (PTP1Btail^N412I^, Fig. 6) localized exclusively to the ER with no detectable presence at the mitochondria (Fig. 13). As the wild-type tail anchor already is restricted in its localization to the ER/vacuole in yeast, we expected that this mutation would have no effect. Indeed, this mutant tail anchor localized identically to the wild-type in yeast (data not shown).

**Figure 13.**
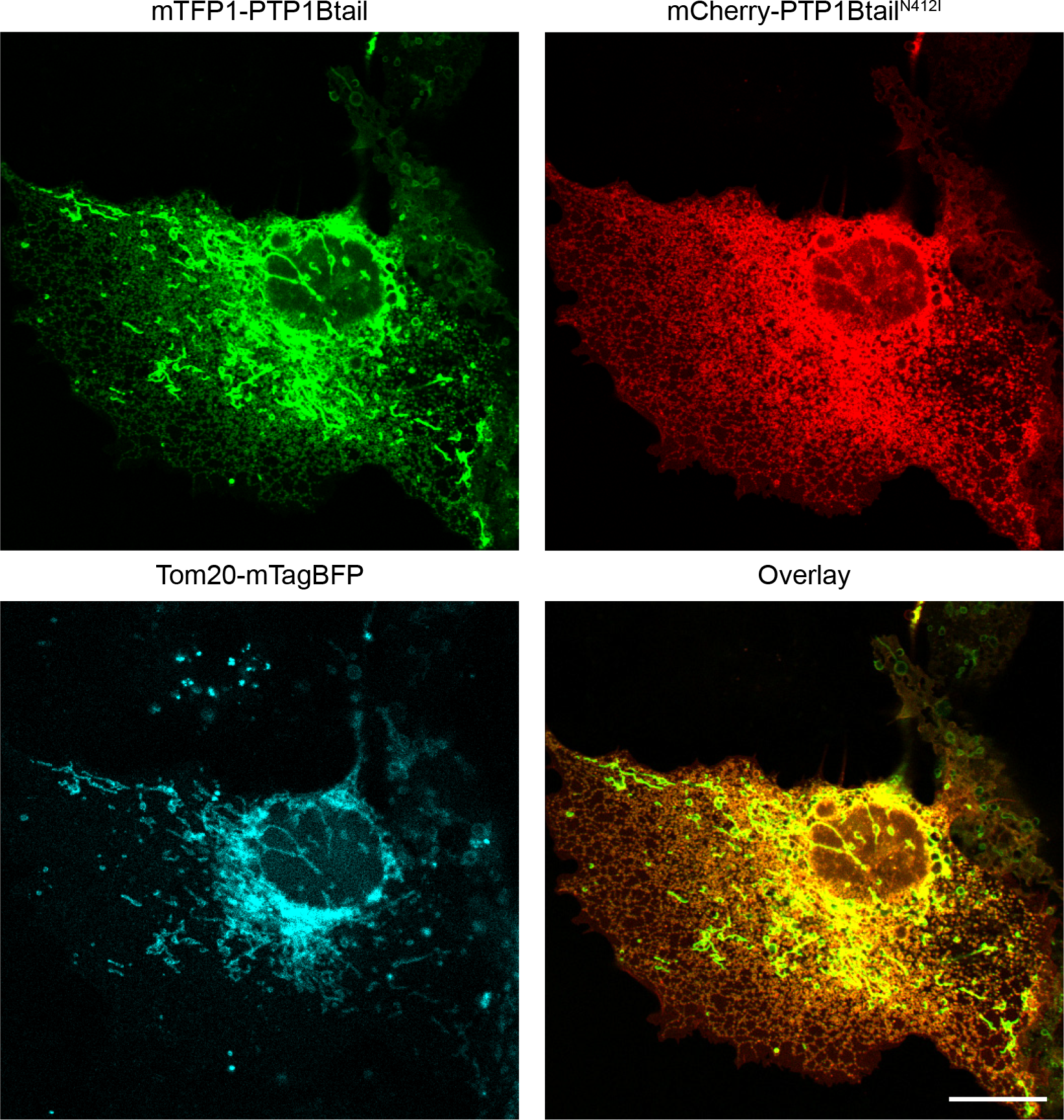
Localization of a tail isoform with high hydropathy in COS-7 cells. The wild-type tail anchor chimera mTFP1-PTP1Btail is shown (green) alongside tail isoform chimera mCherry-PTP1Btail^N412I^ having higher hydropathy (red) in COS-7 cells. Their red/green overlay is also displayed as well as the mitochondrial marker Tom20-mTagBFP. Scale bar: 20 μm.

In the above, we have shown that three generic features of the tail anchor — namely, its TMD length, charge, and hydropathy — appear to largely account for its subcellular targeting (summarized in Fig. 14). To test this further, we constructed a mutant version of the tail anchor for which the TMD was completely scrambled in its amino acid sequence in a way that closely preserved its exact hydropathic profile (by exchanging each original amino acid with a similarly hydrophobic amino acid, Fig. 6). Efficient cloning of this scrambled isoform using long primers required truncating the original tail anchor by four amino acids. We therefore first tested the effect of this mild truncation on subcellular partitioning. As already mentioned above, the terminal 31 amino acids of the tail (PTP1Btail^ΔHALS^, Fig. 6) are sufficient to account for the “wild-type” targeting of PTP1B to the mitochondria and ER (Fig. 15A). A scrambled tail isoform (PTP1Btail^Scr^, Fig. 6) was then constructed that has, by design, a very similar hydropathy profile (Fig. 15A). The scrambled tail localized to the ER and the Golgi, as well as to rapidly moving punctate vesicles, with no discernible presence at the mitochondria (Figs. 15B and 15C). This striking difference in localization from the original PTP1Btail^ΔHALS^ isoform (or, as well, the wild-type anchor) suggests that — in addition to its TMD length, charge, and hydropathy (Fig. 14) — the *exact* amino acid sequence of the tail also appears to contribute to its exact subcellular targeting.

**Figure 14.**
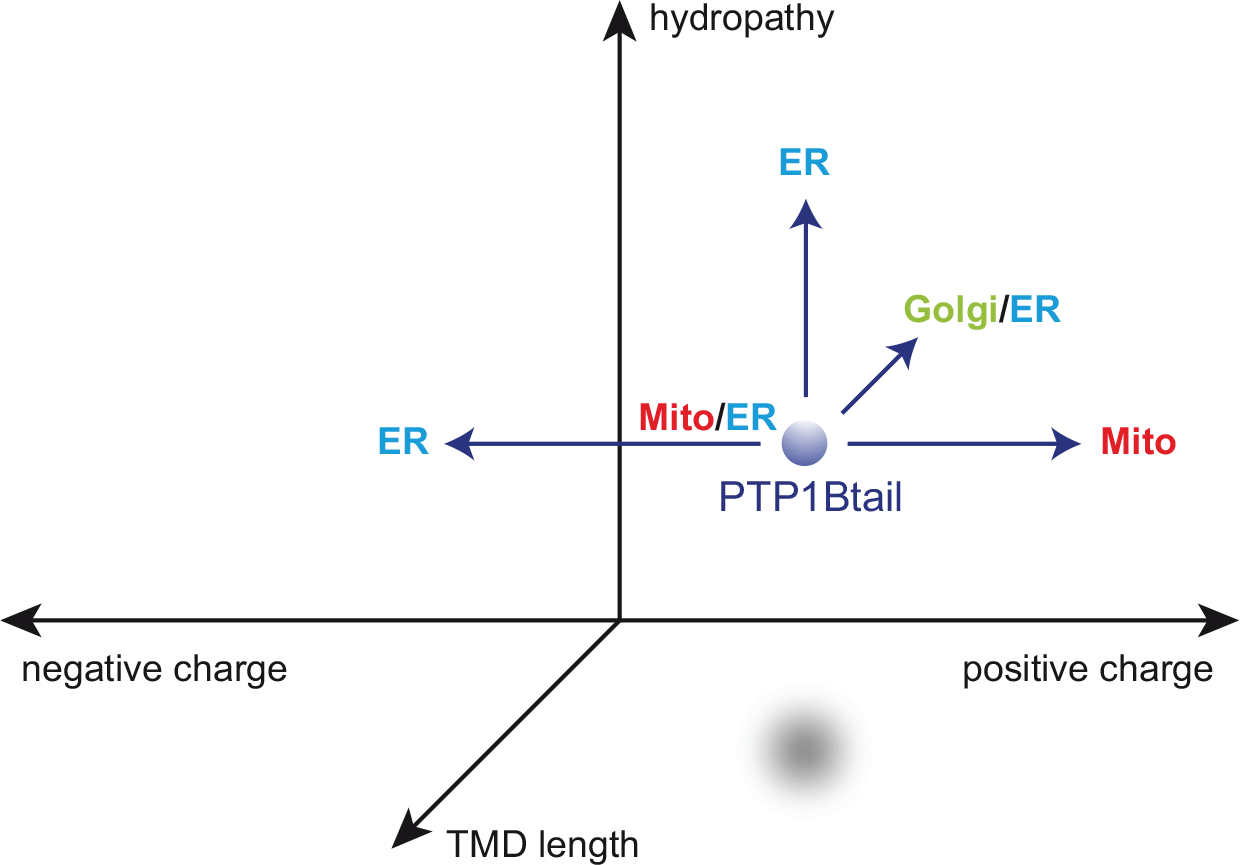
Tail anchor targeting is largely dictated by three different factors: TMD length, C-terminal charge, and hydropathy. Our experimental localization results for different tail isoforms in COS-7 cells are summarized. We have shown that the wild-type localization of PTP1Btail to the mitochondria and ER (Mito/ER) is affected by changes in any of these three factors (see text and Figs. 11–13).

**Figure 15.**
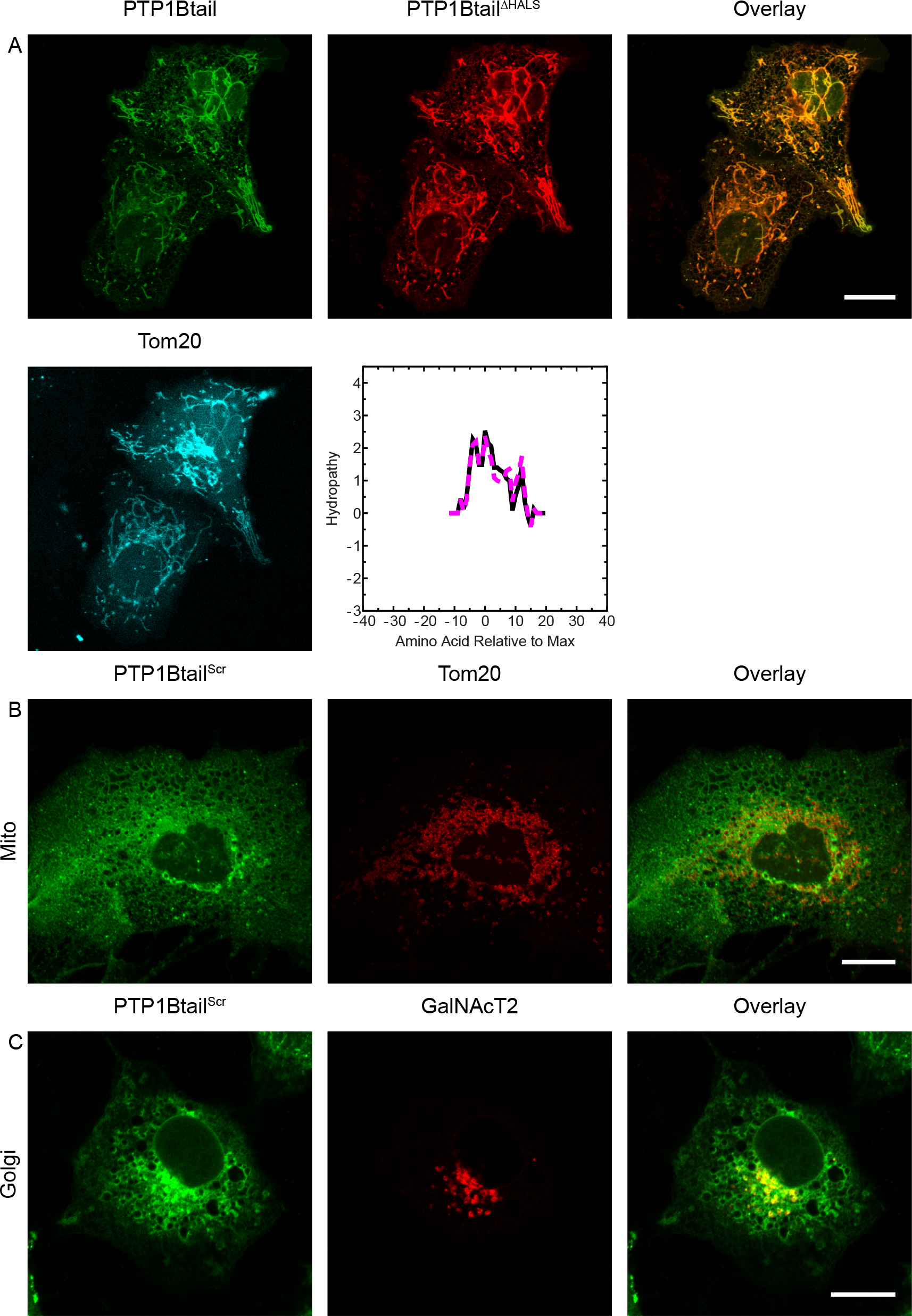
Localization of a scrambled tail isoform in COS-7 cells. (A) Subcellular partitioning of a slightly truncated tail isoform PTP1Btail^ΔHALS^ (Fig. 6, red) and the original tail anchor (green) were identical (see overlay), with both accumulating strongly at the mitochondria (as marked using Tom20-mTagBFP, cyan). The hydropathy profiles of the PTP1Btail^ΔHALS^ (black) and a scrambled isoform PTP1Btail^Scr^ (Fig. 6, dashed magenta) are also shown. (B–C) Coexpression of the fluorophore-labeled PTP1Btail^Scr^ in COS-7 cells (mCherry-PTP1Btail^Scr^) along with either the mitochondrial marker Tom20-mTagBFP (B) or the Golgi marker GalNAcT2-mTurquoise (C). Scale bars: 20 μm.

### Probing the Regulation of Mitochondrial Tyrosine Signaling by PTP1B

Tyrosine phosphorylation of multiple proteins in the mitochondria appears to be a significant regulatory mechanism for general mitochondrial functions^34^. The presence of PTP1B at the mitochondria could be important for the local regulation of these phosphotyrosine-containing targets. To examine the possible presence of tyrosine phosphorylation at the mitochondria, we employed two commonly used probes that recognize phosphotyrosines, one containing a double Src homology 2 domain (dSH2-YFP^75^) and the other containing the phosphotyrosine-binding domain of Shc (PTB-mCherry^76^).

As mentioned above, previous studies have revealed a significant mitochondrial population of the non-receptor tyrosine kinase c-Src as well as other members of the Src family^33,36–40^. To probe Src family activity at the mitochondria, we examined the localization of the dSH2-YFP probe in COS-7 cells. We observed no significant mitochondrial localization of this probe either before or after EGF stimulation (Fig. 16). However, it is likely that this probe can only access the cytosolic face of the OMM, implying at any rate no significant active c-Src (or other family members?) at this submitochondrial region in COS-7 cells.

**Figure 16.**
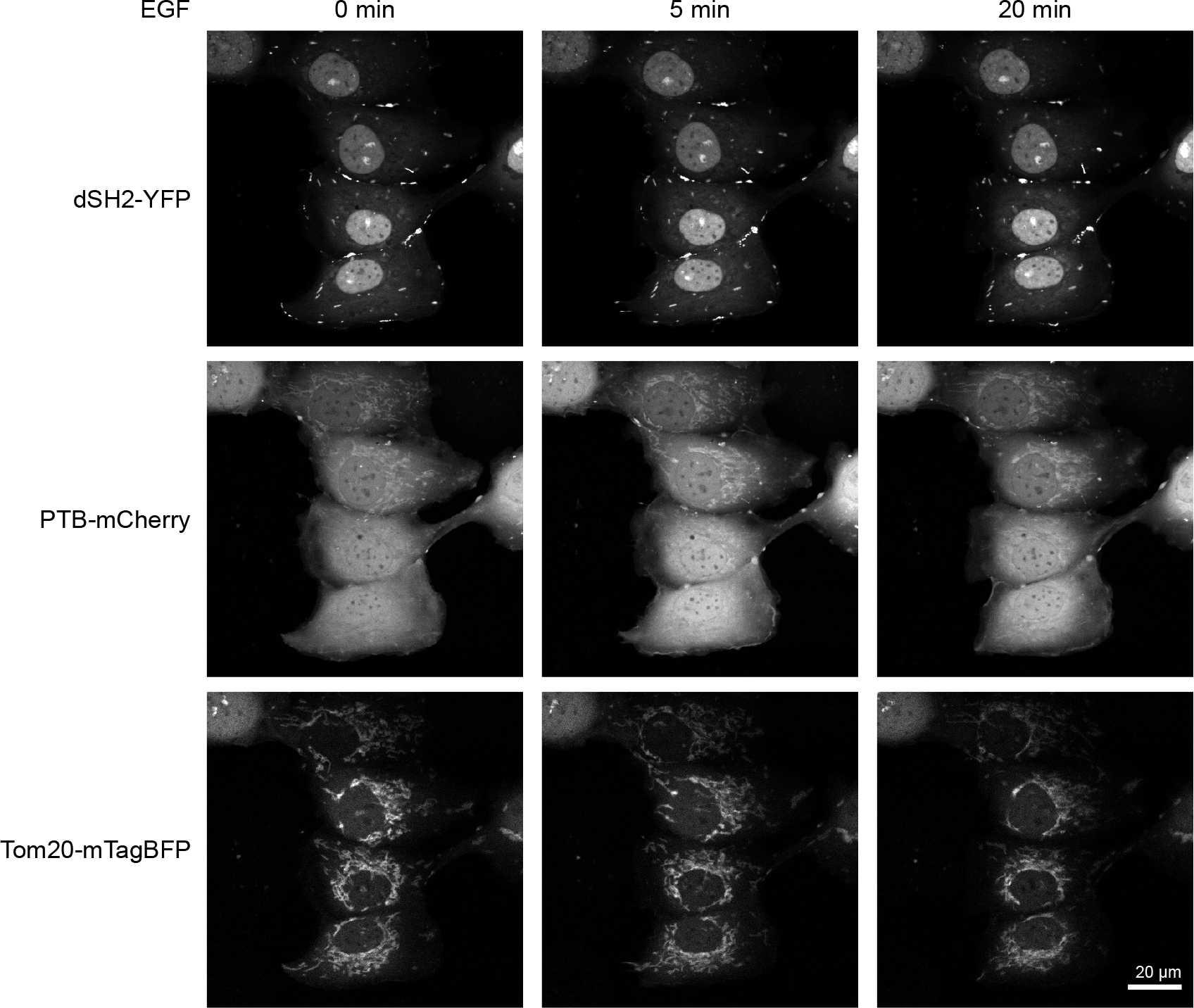
Visualizing mitochondrial phosphotyrosine before and after EGF stimulation. COS-7 cells expressing dSH2-YFP, PTB-mCherry, and the mitochondrial marker Tom20-mTagBFP were starved and imaged before (0 min) and after (5 min and 20 min) their stimulation with EGF (100 ng/mL). No mitochondrial localization of dSH2-YFP was observed. In contrast, a basal accumulation of PTB-mCherry at the mitochondria was observed that increased following EGF stimulation. Scale bar: 20 μm.

Localization of the PTB-mCherry probe in the same COS-7 cells was also examined (Fig. 16). PTB-mCherry displayed a basal recruitment to the mitochondria that significantly increased upon EGF stimulation. The PTB-mCherry probe may be detecting important Shc substrates like the receptor tyrosine kinases ErbB1 and ErbB2, for which mitochondrial pools have recently been claimed^41–43^.

Mitochondrial PTP1B should be able to interact with (and regulate) an hypothesized pool of active mitochondrial ErbB1. To test this, we used confocal time-domain FLIM to visualize the direct interaction of donor-labeled ErbB1 (ErbB1-mCitrine) with an acceptor-labeled D181A trapping mutant^11^ of PTP1B (mCherry-PTP1B^D/A^) by FRET. COS-7 cells expressing these constructs were starved overnight and stimulated with EGF. Continuous recording of the donor fluorescence before and during the stimulation allowed dynamic monitoring of the ErbB1-PTP1B interaction across the cell, including at the mitochondria (Fig. 17 and Movie 2). In the top two rows of Fig. 17, cells expressing only ErbB1-mCitrine are shown. These cells exhibited a stable and spatially uniform lifetime of 3.02 ns (the isolated spots of low lifetime are consistent with autofluorescent particles, which were also often observed in untransfected cells). Separate labeling of mitochondria using Tom20-mTagBFP allowed identification of regions of the image containing mitochondria. Surprisingly, ErbB1 was not observed to accumulate to the mitochondria after EGF stimulation (see further discussion below). In the third and fourth rows of Fig. 17, COS-7 cells coexpressing donor-labeled ErbB1 and the acceptor-labeled trapping mutant (mCherry-PTP1B^D/A^) exhibited a low basal interaction before stimulation that was sharply increased upon the addition of EGF. The full lifetime movie of these cells (Movie 2) was consistent with previous claims of the restricted interaction of PTP1B with ErbB1 only after receptor internalization by endocytosis^28^. A high level of FRET-based interaction of ErbB1-mCitrine with mCherry-PTP1B^D/A^ was detectable on perinuclear structures that were consistent with the mitochondria (as visualized with Tom20-mTagBFP). These results suggest an important role for PTP1B in the local dephosphorylation of ErbB1 at the mitochondria, both before and after EGF stimulation.

**Figure 17.**
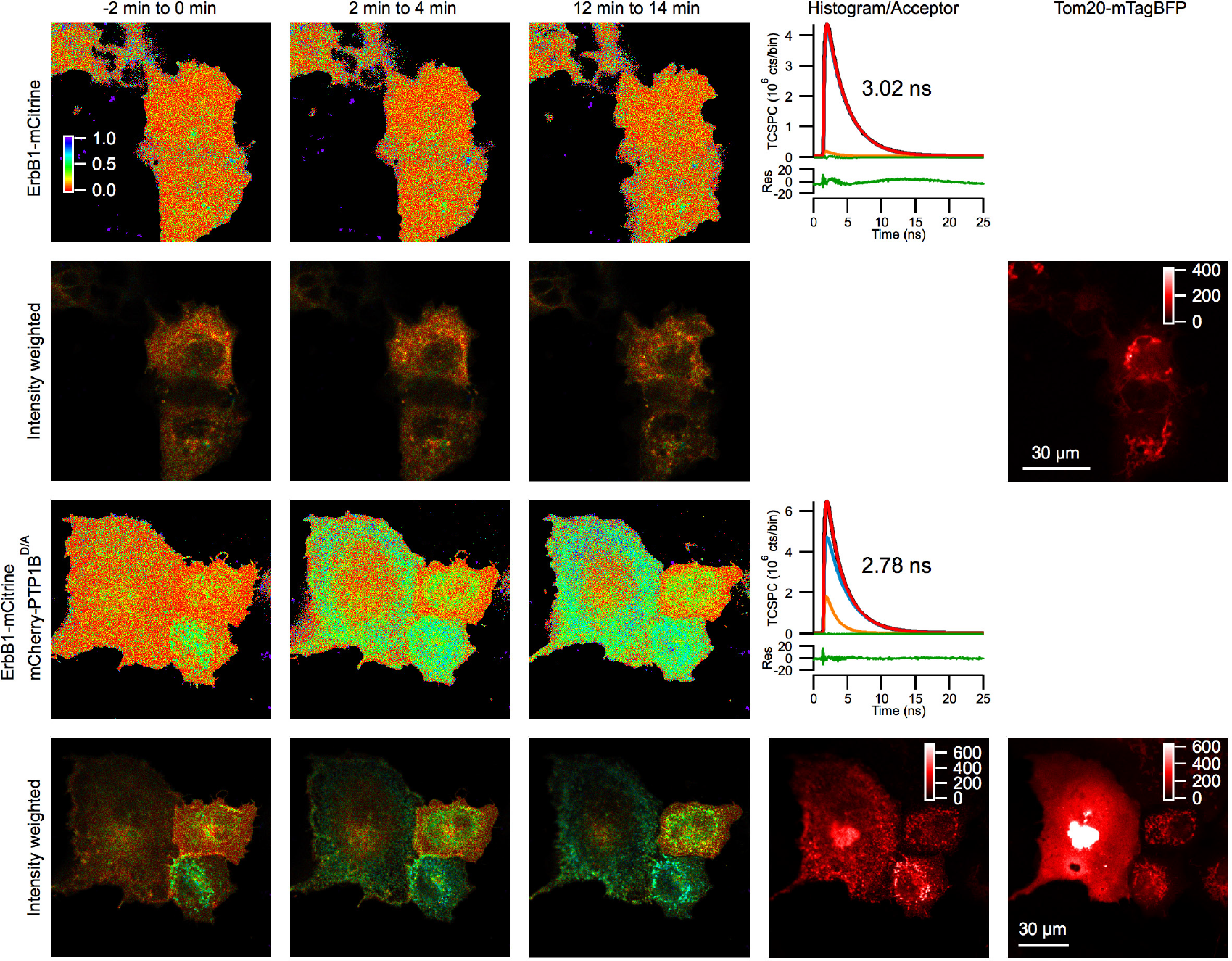
Dynamic FLIM-based monitoring of the subcellular interaction of mCitrine-ErbB1 with mCherry-PTP1B^D/A^ in COS-7 cells following EGF stimulation. Donor lifetime images of mCitrine-ErbB1 are shown before (−2 min to 0 min) and after (2 min to 4 min, 12 min to 14 min) stimulation with EGF in cells expressing the donor-labeled ErbB1 and the mitochondrial marker Tom20-mTagBFP in the first and second rows (lifetime control cells, representative of n=3 recordings). A lifetime of 3.02 ns was obtained from fitting the histogram of the entire 16-minute recording, assuming a fixed donor-only lifetime of 3.05 ns (blue TCSPC histogram) and a FRET lifetime of 1.51 ns (orange TCSPC histogram), which were determined from double lifetime fitting of the cells shown in the third and fourth rows. A generally low FRET fraction (consistent with zero) was obtained across the images. FLIM images of COS-7 cells expressing mCitrine-ErbB1, mCherry-PTP1B^D/A^, and Tom20-mTagBFP are shown in the third and fourth rows (representative of n=5 recordings). Here, a basal interaction (of varying strength) is detected that increases in all of the cells after EGF stimulation. An average lifetime of 2.78 ns over the entire 16-minute movie was obtained, which was significantly lower than the 3.02 ns lifetime of the donor-only control cells. A specific decrease in lifetime at the mitochondria was clearly observed in the cells on the right. The Tom20-mTagBFP mislocalized in the cell on the left (large aggregate) preventing determination of the mitochondria in this cell. Color scale in the top left image gives the FRET fraction α. Scale bars: 30 μm.

As an important control, we also examined cells coexpressing ErbB1-mCitrine with an acceptor-labeled chimera containing only the tail anchor of PTP1B, mCherry-PTP1Btail (Fig. 18). In these cells, no basal interaction was detected and no significant increase in interaction was observed following EGF stimulation. This importantly shows that overexpression of the acceptor-labeled tail is insufficient to affect the lifetime of ErbB1-mCitrine on the ER or the mitochondria. The decreased lifetime in Fig. 17 for the acceptor-labeled trapping mutant is therefore indicative of a direct interaction of ErbB1-mCitrine with mCherry-PTP1B^D/A^.

**Figure 18.**
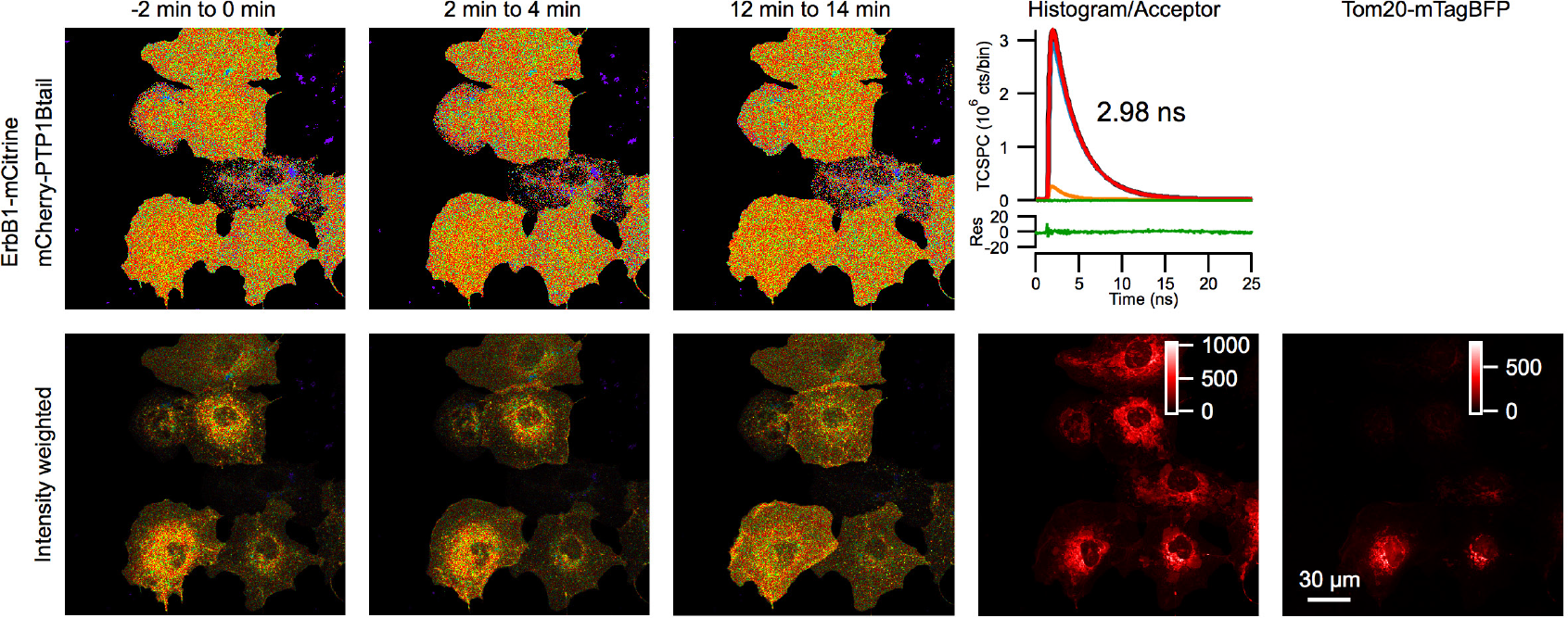
Control for dynamic FLIM-based monitoring of subcellular ErbB1 interaction with PTP1B^D/A^. Donor lifetime images of COS-7 cells expressing ErbB1-mCitrine (donor), mCherry-PTP1Btail (acceptor), and the mitochondrial marker Tom20-mTagBFP (representative of n=3 recordings, see Fig. 17 for further details). The average lifetime of the entire 16 minute recording was 2.98 ns. A generally low FRET fraction α across the cells was detected that was similar to the negative control shown in the first two rows of Fig. 17. Despite the similar localization and expression of the tail-only acceptor-labeled chimera, no lifetime reduction either before or after EGF stimulation was detectable of the donor-labeled mCitrine-ErbB1. This control, therefore, importantly demonstrates that the reduced lifetime in the bottom two rows of Fig. 17 reflected the direct interaction of mCitrine-ErbB1 with the catalytic domain of mCherry-labeled PTP1B^D/A^. Scale bar: 30 μm.

To better distinguish the interaction of ErbB1 with PTP1B^D/A^ across the cell, we also performed similar experiments with different chimeras that localized the catalytic domain of the trapping mutant exclusively to the ER (C-terminal fusion of the tail anchor of the Sec61β subunit^25^, mCherry-PTP1B^D/A^-ER, first and second rows of Fig. 19) or to the OMM (C-terminal fusion of the +2 positively charged mutant tail anchor of CytB5 that localizes exclusively to the OMM^73^, mCherry-PTP1B^D/A^-OMM, third and fourth rows of Fig. 19). For both the ER- and OMM-localizing chimeras, a low basal interaction before stimulation and a robust increase in interaction after stimulation were observed only at these respective compartments. These results importantly reveal that the interaction of PTP1B^D/A^ with phosphorylated ErbB1 involves direct interaction of these proteins at the ER or the cytosolic face of the OMM.

**Figure 19.**
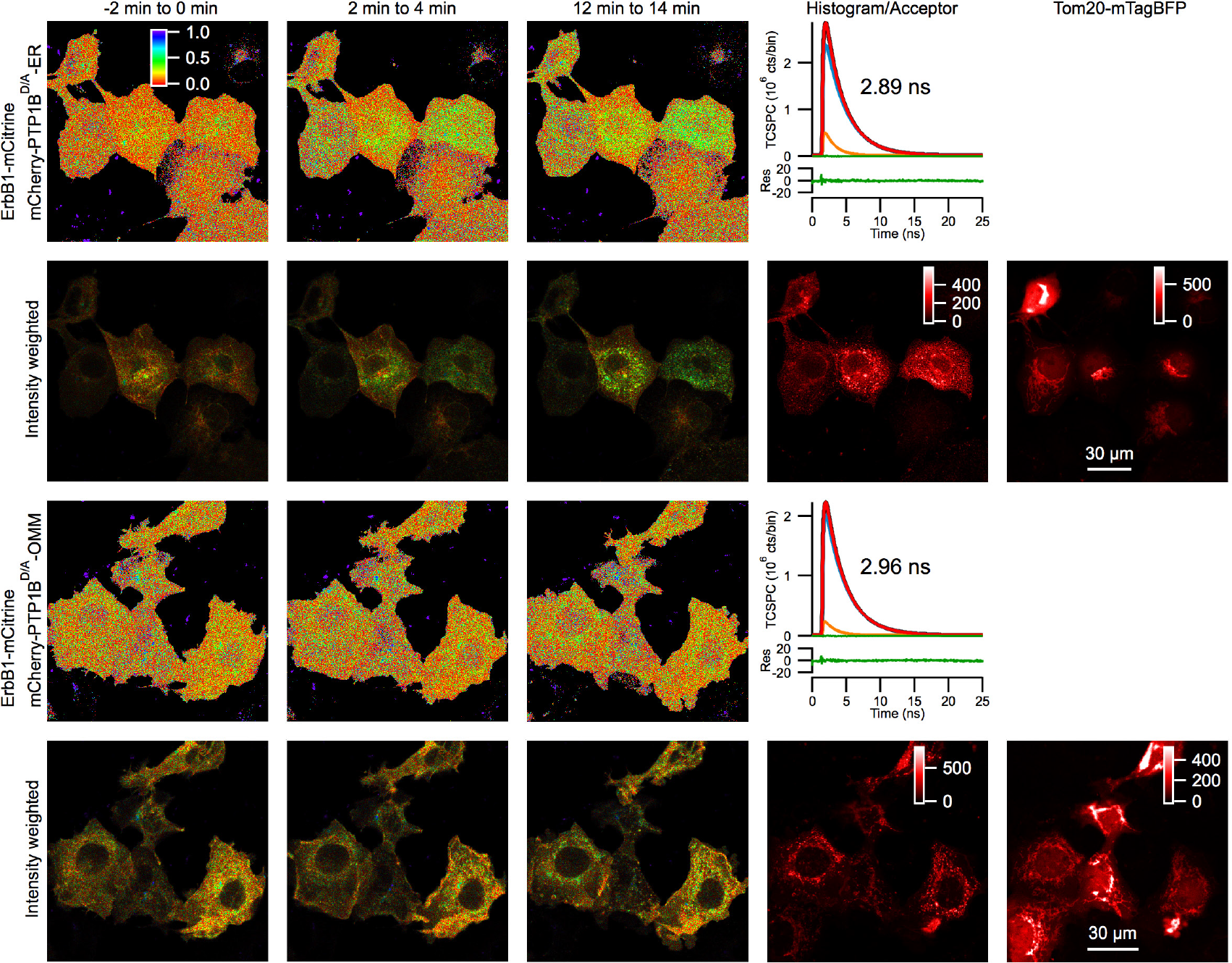
Dynamic FLIM-based monitoring of the interaction of mCitrine-ErbB1 with mCherry-labeled PTP1B^D/A^ targeted to either the ER or the outer mitochondrial membrane. In the first two rows, donor lifetime images of COS-7 cells expressing ErbB1-mCitrine, mCherry-PTP1B^D/A^-ER, and the mitochondrial marker Tom20-mTagBFP are displayed before and after EGF stimulation (representative of n=3 recordings, see Fig. 17 for further details). A robust decrease in lifetime was detectable upon EGF stimulation, revealing the specific interaction of ErbB1 with ER-localized PTP1B. In the third and fourth rows, donor lifetime images of COS-7 cells expressing ErbB1-mCitrine, mCherry-PTP1B^D/A^-OMM, and the mitochondrial marker Tom20-mTagBFP are displayed before and after EGF stimulation (representative of n=4 recordings). A robust decrease in lifetime was detected upon EGF stimulation only in the vicinity of the mitochondria (the mCherry-PTP1B^D/A^-OMM construct and the mitochondrial marker Tom20-mTagBFP overlapped well). Despite the only slight reduction of the lifetime of the entire 16 minute recording (2.89 ns for the first two rows, 2.96 ns for the last two rows), local lifetime reductions were clearly detected that coincided with the acceptor localization (ER in top two rows, mitochondria in bottom two rows). Scale bars: 30 μm.

We next attempted to resolve whether a direct interaction between ErbB1 and PTP1B^D/A^ could be detected *within* the mitochondria. We therefore examined the interaction of donor-labeled ErbB1 with an acceptor-labeled PTP1B^D/A^ chimeras targeted either to the IMS (N-terminal fusion of Smac domain^77^, mCherry-PTP1B^D/A^-IMS, first and second rows of Fig. 20) or to the matrix (N-terminal fusion of COX8A domain^78^, mCherry-PTP1B^D/A^-MAT, third and fourth rows of Fig. 20). In neither case were we able to detect a decreased lifetime at the mitochondria. As for the control cells shown in the first two rows of Fig. 17, no significant concentration of ErbB1 could be detected at the mitochondria before or after EGF stimulation.

**Figure 20.**
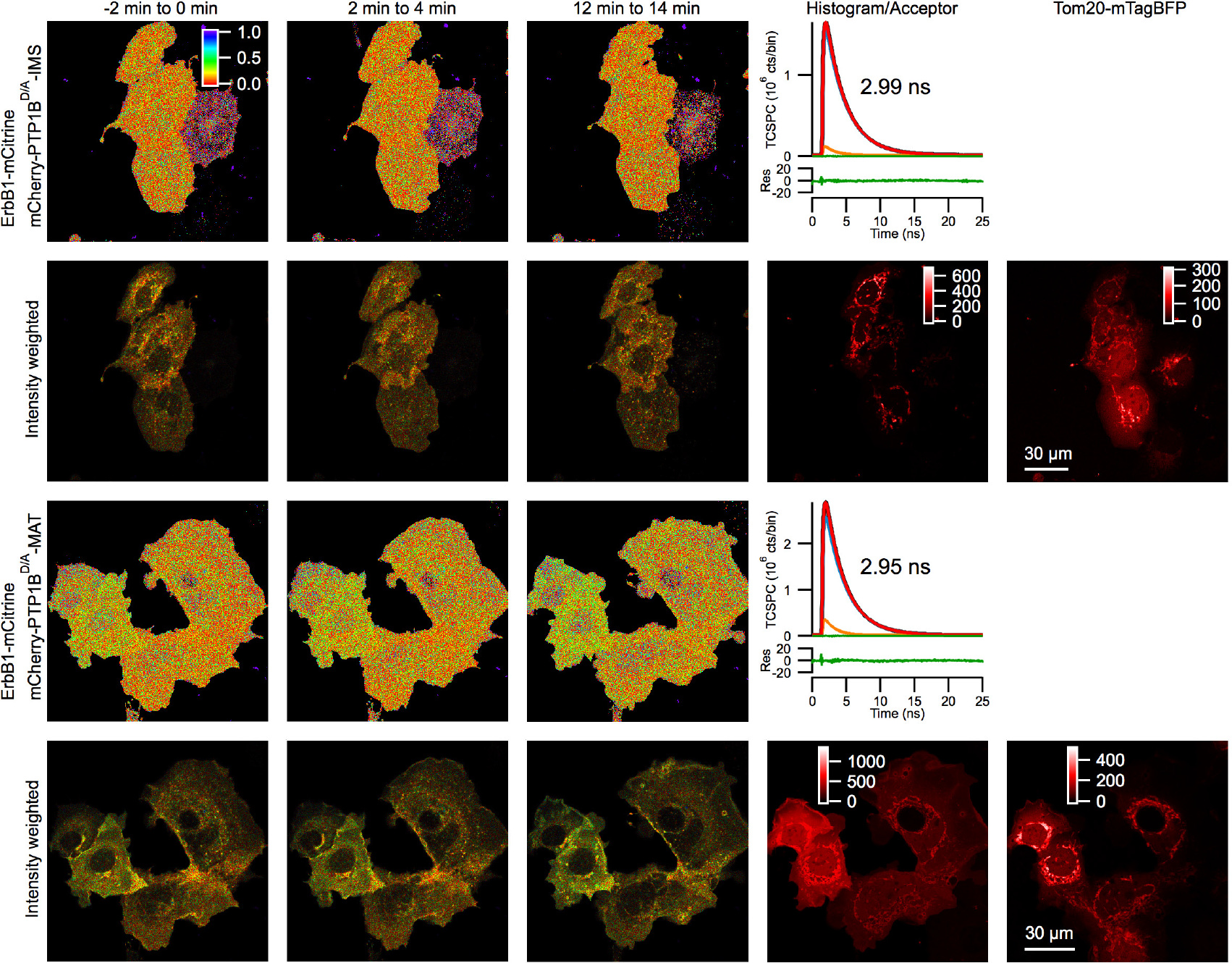
Dynamic FLIM-based monitoring of the interaction of mCitrine-ErbB1 with mCherry-labeled PTP1B^D/A^ targeted to either the intermembrane space (IMS) or the mitochondrial matrix (MAT). In the first two rows, donor lifetime images of COS-7 cells expressing ErbB1-mCitrine, mCherry-PTP1B^D/A^-IMS, and the mitochondrial marker Tom20-mTagBFP are displayed before and after EGF stimulation (representative of n=8 recordings, see Fig. 17 for further details). No significant decrease in lifetime was detectable upon EGF stimulation, either generally across the cell or specifically at the mitochondria. In the third and fourth rows, donor lifetime images of COS-7 cells expressing ErbB1-mCitrine, mCherry-PTP1B^D/A^-MAT, and the mitochondrial marker Tom20-mTagBFP are displayed before and after EGF stimulation (representative of n=8 recordings). While there was a generally decreased lifetime in some cells (reflected by the lower lifetime of 2.95 ns obtained from fitting the entire 16 minute recording), no specific decrease (either basal or time-dependent) was detectable at the mitochondria. Scale bars: 30 μm.

As previous studies reporting the localization of ErbB1 or ErbB2 to mitochondria were performed using cells that either naturally overexpressed c-Src (breast cancer cells^43^) or were cotransfected with a c-Src construct (10T1/2 cells^41,42^), we undertook a separate set of experiments in which COS-7 cells were transfected with ErbB1-mCitrine, c-Src-mTurquoise, and Tom20-mTagBFP. In these experiments, we also observed no detectable mitochondrial presence of the fluorescently labeled c-Src or ErbB1 (data not shown). We also performed a set of experiments using MCF-7 cells, which naturally express high levels of c-Src, but again we were unable to see mitochondrial recruitment of fluorescent ErbB1 either before or after EGF treatment (in cells doubly transfected with ErbB1-mCitrine and Tom20-mTagBFP, data not shown). The discrepancy of our current experiments with previous reports of ErbB1 (and c-Src) at the mitochondria could be due to a number of factors (cell line specificity, an only small fraction of mitochondrial-localizing ErbB1/c-Src, or more detailed experimental differences between our live cell studies and previous studies based on fixed cells and mitochondrial extraction^41,42^).

While we were unable to visualize the independent recruitment of ErbB1 to the mitochondria, our FLIM experiments nevertheless collectively demonstrate a robust interaction of mCherry-PTP1B^D/A^ with ErbB1-mCitrine at the OMM. This importantly reveals that PTP1B has a significant presence at the OMM (which was unclear based only on our electron micrographs, Fig. 5), where it likely can engage intracellular ErbB1.

## Discussion

We have shown that the tail anchor of PTP1B, while traditionally considered to target it only to the ER, actually guides it efficiently to the mitochondria in mammalian cells (using fluorescence-based detection of endogenous/overexpressed PTP1B), with clear evidence for an accumulation of PTP1B at MAM sites along the ER (revealed by electron microscopy), a significant pool at the OMM (from our FLIM studies), and a high concentration of PTP1B distributed throughout the mitochondrial matrix (electron microscopy). Our studies in yeast point to spontaneous insertion as the likely dominant mechanism for its entry into the ER membrane. Its accumulation at MAM sites along the ER suggests a possible role for these regions in facilitating its transfer from the ER to the mitochondria; however, spontaneous entry as well into the mitochondrial membrane cannot be ruled out. While our studies have collectively reduced the spectrum of possibilities, the *exact* mechanisms (whether spontaneous or protein-mediated) that ultimately control PTP1B’s tail-anchor-driven insertion into the ER, its accumulation at MAM sites, its insertion into the OMM, and its entry into the mitochondrial matrix require still further exploration.

To better understand how PTP1B’s subcellular partitioning is determined by its tail anchor, we undertook a systematic exploration of the tail anchor and related isoforms in both mammalian cells and yeast. Our investigations in mammalian cells and yeast revealed important information with regard to (1) the properties of the tail anchor itself (TMD length, charge, and hydropathy), (2) the properties of the subcellular organelles of the host cell (e.g. lipid composition), and (3) differences in the properties of a given organelle for widely evolutionarily separated host cells. These explorations yielded many surprising results on PTP1B’s tail-anchor localization (summarized in Table 1).

**Table 1.**
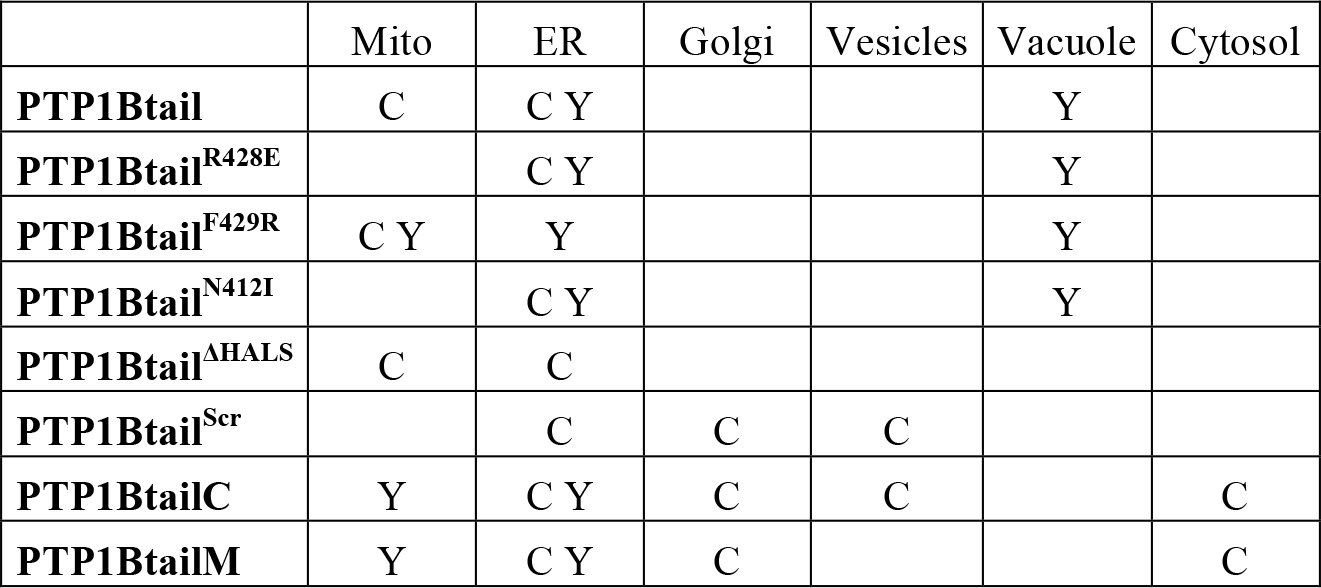
Localization in COS-7 (C) and yeast (Y) of different fluorophore-labeled PTP1B constructs.

Upon heterologous expression of the tail anchor in yeast, we found that it localized to the ER and vacuole and not at all to the mitochondria (Fig. 8) despite its similar hydropathy as compared with mitochondrial tail-anchor proteins in yeast (Fig. 7). This was in line with previous observations of the heterologous expression in yeast of the mammalian protein Bcl2, which localizes to the ER/mitochondria in mammalian cells^79,80^ but only to the ER in yeast^81^.

We found no difference of the localization of the tail anchor in yeast strains that synthesized ergosterol versus cholesterol (Fig. 9). However, the *overall* sterol concentration (or its concentration within rafts) might still affect global (or local) membrane thickness and fluidity for specific organelles in mammalian cells versus yeast^70^. Previous in vitro studies have demonstrated that membranes with high sterol content inhibit the spontaneous insertion of the tail anchor proteins CytB5^82^ and PTP1B^27^, suggesting that overall sterol concentration might still contribute to the observed subcellular partitioning of PTP1B in mammalian cells and yeast.

While a slight N-terminal truncation of the 35 amino acid tail anchor to 31 amino acids preserved its localization, more drastic truncations from the N- and/or C-termini localized to the ER/Golgi in mammalian cells and to the ER/mitochondria in yeast. The collective differences of the wild-type (discussed above) and the highly truncated isoforms upon expression in mammalian cells versus yeast likely reveal important differences in tail anchor targeting across the evolutionary tree. These observations may reflect important differences in the properties of the diverse organelles with a given host cell and for a specific organelle across the evolutionary tree. As already discussed above, the specific lipid composition of the membrane of an organelle can, for example, affect bilayer width, charge, membrane fluidity, and curvature and therefore the insertion stability of a given tail anchor.

Regarding C-terminal tail anchor charge, the addition of a positive charge to the C-terminus shifted PTP1B completely (mammalian cells) or largely (yeast) to the mitochondria. This positive-charge-based relocation to the mitochondria is likely attributable to the presence of higher net negative charge in these compartments (the negatively charged mitochondrial-specific lipid cardiolipin^83,84^). Too little positive charge in the wild-type tail anchor of PTP1B appears to be the most probable explanation for accounting for its absence from yeast mitochondria, indicating that yeast mitochondrial membranes have an overall lower negative charge than mammalian mitochondria.

Our results on TMD length and C-terminal charge, along with our own current results (Fig. 13) as well as prior results (mutants of the Fis1 tail anchor^85^) on hydropathy, contribute to the emerging view that these three properties play an important role in specifying the subcellular targeting of a given tail anchor. However, the exact amino acid sequence appears to also play a role (see our results on the scrambled tail anchor in Fig. 15).

Similar systematic exploration of other tail anchors should help confirm the importance of the effect of the three dimensions of TMD length, charge, and hydropathy (see Fig. 14) on subcellular partitioning. Tail anchor isoforms that reside in certain regions of this 3D phase space may act as “restricted keys” for accessing the lipid bilayer of only a single compartment (e.g. mitochondria alone), whereas tail anchors residing in other regions may act as “skeleton keys” permitting access to multiple compartmental membranes (e.g. ER/mitochondria or ER/Golgi). Design principles based on the tail anchor properties outlined above could be used to construct synthetic tails with high specificity for each organellar membrane in the cell^86,87^. Further examination of these aspects of tail anchor targeting, in addition to a more detailed portrait of the global and local concentrations of particular lipid isoforms (including charged isoforms and isoforms that can affect bilayer width), should help to account for the observed partitioning of general tail anchor proteins in both higher and lower eukaryotes. While this simple three-dimensional view of TMD length, charge, and hydropathy is compelling, other sequence-specific aspects should not be ignored, as we have demonstrated through hydropathy-preserving scrambling of the amino acids in the TMD (Fig. 15).

PTP1B’s tail anchor-dependent presence at and within the mitochondria likely reflects its important physiological roles in the regulation of phosphotyrosine-based signaling at the outer membrane (e.g. regulation of mitochondrial EGFR signaling^41–43)^ and in the mitochondrial interior (e.g. regulation of activities of Src^33,36–40^ and SHP2^33,40,44^, regulation of electron transport chain enzymes^34,35^). Further investigations of PTP1B’s interactions with known and putative substrates using standard biochemical approaches or advanced microscopy would be useful. In particular, our library of PTP1B^D/A^ chimeras that localize to each submitochondrial region (OMM, IMS, matrix) should provide a useful set of tools for future FLIM-based examination of its locally restricted engagement with substrates. Its potential regulation of basic mitochondrial functioning could be revealed using assays for cellular oxygen consumption, electron transport chain activity, glucose uptake, lactate production, or the ATP/ADP ratio^43^. Finally, methods for detecting different post-translationally modified isoforms of PTP1B (phosphorylated, oxidized, sumoylated, cleaved) would be useful to develop and employ for discerning any potential differences (that may have functional consequences) of mitochondrial PTP1B subfractions from its more distributed pool along the ER.

Such deeper investigations of the role of this newly identified pool of mitochondrial PTP1B on the regulation of local tyrosine-based signaling and potentially fundamental mitochondrial processes should help shed light on its apparently complex roles in normal and diseased states^6,7^. Moreover, our use of targeted mutagenesis combined with heterologous expression in yeast revealed intriguing and likely fundamental differences in tail-anchor targeting to different organelles within a single host cell and for targeting to the same organelle in widely separated hosts across the evolutionary tree.

## Materials and Methods

### Yeast plasmids

The constructs pAK51 (yemCitrine-PTP1Btail), pAK72 (yemCitrine-Ysy6tail), and pAK86 (yemCitrine-Sec22tail) that we used for construction of the respective strains sAK199, sAK238, and sAK264 (Table 2) were derived from pYM-N17 (see Janke et al.^88^). Briefly, the resistance cassette containing the gene for resistance to ClonNAT was reversed between the SalI/SacI sites of pYM-N17 to generate pYM-N17-Natrev. Through a series of changes in the latter construct, we obtained pAK51, in which the GPD promoter is followed by CGGATTCTAG<u>GCTAGC</u>CGCCGCC (unique NheI site underlined), the sequence for yemCitrine without its stop codon (gift from the Knop lab), a short linker, the sequence for the C-terminal 35 amino acids of PTP1B (tail anchor), followed by a unique EcoRI site, and then the Tcyc1 terminator (with its insertion removing the original EcoRI site from pYM-N17-Natrev). To obtain pAK72 (or pAK86), an NheI-containing 5′ primer of yemCitrine was used along with a long EcoRI-containing 3′ primer of yemCitrine containing the C-terminal 31 amino acids of Ysy6 (or C-terminal 31 amino acids of Sec22) was used to generate the PCR product NheI-yemCitrine-Ysy6-EcoRI (or NheI-yemCitrine-Sec22-EcoRI) for replacement of yemCitrine-PTP1Btail between the NheI and EcoRI sites of pAK51. Plasmids pAK87 (yemCitrine-PTP1BtailM) and pAK88 (yemCitrine-PTP1BtailC), which contained the amino acid sequences given in Fig. 6, were constructed in a similar fashion. Plasmid pAK104 (yemCitrine-PTP1Btail^R428E^) was constructed by double point mutation of the codon for arginine in pAK51 (AGG→GAG) as well as pAK111 (yemCitrine-PTP1Btail^F429R^) by double point mutation of the indicated codon for phenylalanine (TTC→CGC). Plasmid pAK109 (yemCitrine-PTP1Btail^N412I^) was constructed by point mutation of the indicated codon for asparagine (AAC→ATC).

**Table 2.** Yeast strains. All strains with an AK prefix — and therefore not ESM356-1^91^, RH2881^59^, and RH6829^59^ — were generated for this study.

| strain |  |
| --- | --- |
| ESM356-1 | MATa ura3-52 leu2Δ1 trp1Δ63 his3Δ200 |
| AK199 | ESM356-1 leu2Δ0::NatNT2rev-GPD-yemCitrine-PTP1Btail-Tcyc1 |
| AK201 | ESM356-1 leu2Δ0::NatNT2rev-GPD-yemCitrine-PTP1Btail-Tcyc1 STE2::mCherry::kanMX |
| AK202 | ESM356-1 leu2Δ0::NatNT2rev-GPD-yemCitrine-PTP1Btail-Tcyc1 COX4::mCherry::kanMX |
| AK203 | ESM356-1 leu2Δ0::NatNT2rev-GPD-yemCitrine-PTP1Btail-Tcyc1 SEC7::mCherry::kanMX |
| AK204 | ESM356-1 leu2Δ0::NatNT2rev-GPD-yemCitrine-PTP1Btail-Tcyc1 CWP2::mCherry::kanMX |
| AK205 | ESM356-1 leu2Δ0::NatNT2rev-GPD-yemCitrine-PTP1Btail-Tcyc1 get3Δ::klURA |
| AK206 | ESM356-1 leu2Δ0::NatNT2rev-GPD-yemCitrine-PTP1Btail-Tcyc1 get2Δ::klURA |
| AK207 | ESM356-1 leu2Δ0::NatNT2rev-GPD-yemCitrine-PTP1Btail-Tcyc1 sgt2Δ::klURA |
| AK222 | ESM356-1 leu2Δ0::NatNT2rev-GPD-yemCitrine-PTP1Btail-Tcyc1 get2Δ::klURA get1Δ::HIS3MX6 |
| AK225 | ESM356-1 leu2Δ0::NatNT2rev-GPD-yemCitrine-PTP1Btail-Tcyc1 srp101Δ::klURA |
| AK238 | ESM356-1 leu2Δ0::NatNT2rev-GPD-yemCitrine-YSY6tail-Tcyc1 |
| AK256 | ESM356-1 leu2Δ0::NatNT2rev-GPD-yemCitrine-YSY6tail-Tcyc1 get2Δ::klURA |
| AK257 | ESM356-1 leu2Δ0::NatNT2rev-GPD-yemCitrine-PTP1BtailM-Tcyc1 |
| AK258 | ESM356-1 leu2Δ0::NatNT2rev-GPD-yemCitrine-PTP1BtailC-Tcyc1 |
| AK259 | ESM356-1 leu2Δ0::NatNT2rev-GPD-yemCitrine-PTP1Btail-Tcyc1 ydj1Δ::klURA |
| AK260 | ESM356-1 leu2Δ0::NatNT2rev-GPD-yemCitrine-PTP1Btail-Tcyc1 ram1Δ::klURA |
| AK261 | ESM356-1 leu2Δ0::NatNT2rev-GPD-yemCitrine-PTP1Btail-Tcyc1 apj1Δ::klURA |
| AK263 | ESM356-1 leu2Δ0::NatNT2rev-GPD-yemCitrine-PTP1Btail-Tcyc1 sec72Δ::klURA |
| AK264 | ESM356-1 leu2Δ0::NatNT2rev-GPD-yemCitrine-SEC22tail-Tcyc1 |
| AK265 | ESM356-1 leu2Δ0::NatNT2rev-GPD-yemCitrine-SEC22tail-Tcyc1 get2Δ::klURA |
| AK266 | ESM356-1 leu2Δ0::NatNT2rev-GPD-yemCitrine-PTP1Btail-Tcyc1 get2Δ::klURA get1Δ::HIS3MX6 sec72Δ::hphNT1 |
| AK273 | ESM356-1 leu2Δ0::NatNT2rev-GPD-yemCitrine-PTP1BtailM-Tcyc1 SEC7::mCherry::kanMX |
| AK274 | ESM356-1 leu2Δ0::NatNT2rev-GPD-yemCitrine-PTP1BtailC-Tcyc1 SEC7::mCherry::kanMX |
| RH2881 | MATa ura3 leu2 his3 trp1 can1 bar1 |
| AK275 | RH2881 p415-yemCitrine-PTP1Btail |
| RH6829 | MATa ura3 leu2 his3 trp1 can1 bar1 erg5Δ::HIS5-TDH3-DHCR24 erg6Δ::TRP1-TDH3-DHCR7 |
| AK276 | RH6829 p415-yemCitrine-PTP1Btail |
| AK277 | ESM356-1 leu2Δ0::NatNT2rev-GPD-yemCitrine-PTP1BtailR428E-Tcyc1 |
| AK280 | ESM356-1 leu2Δ0::NatNT2rev-GPD-yemCitrine-PTP1BtailN412I-Tcyc1 |
| AK281 | ESM356-1 leu2Δ0::NatNT2rev-GPD-yemCitrine-PTP1BtailF429R-Tcyc1 |
| AK282 | ESM356-1 leu2Δ0::NatNT2rev-GPD-yemCitrine-PTP1BtailR428E-Tcyc1 COX4::mCherry::kanMX |
| AK284 | ESM356-1 leu2Δ0::NatNT2rev-GPD-yemCitrine-PTP1BtailF429R-Tcyc1 COX4::mCherry::kanMX |

The expression plasmid pAK100 (p415-yemCitrine-PTP1Btail) was constructed by PCR of yemCitrine-PTP1Btail from pAK51 flanked by SalI and XhoI restriction sites for insertion into SalI/XhoI-cut p415-GPD-mCherry (gift from M. Knop).

### Mammalian plasmids

All mammalian plasmids listed below utilized pcDNA3.1(+/−) backbones (Clontech) with the indicated genes fused with standard genetically encoded fluorophores (see Walther et al.^89^ for citations) and expressed using a cytomegalovirus (CMV) promoter.

The following plasmids used in this study were: mCitrine-PTP1B (described in Yudushkin et al.^29^), Tom20-mTagBFP and mTagBFP-Sec61β (gifts of R. Stricker and E. Zamir), GalNAcT2-mTurquoise (gift of K. van Eickels), mTurquoise-PTP1B and ErbB1-mTurquoise (gifts of J. Luig), YFP-dSH2 (two consecutive phosphotyrosine-binding Src-homology 2 domains derived from pp60^c-Src^ described in Kirchner et al.^75^), PTB-mCherry (mCherry version of PTB-YFP described in Offterdinger et al.^76^, gift of J. Ibach and P. Verveer), ErbB1-mCitrine (monomeric version of ErbB1-Citrine described in Offterdinger & Bastiaens^90^, gift of J. Ibach and P. Verveer), and mCherry-PTP1B^D/A^ (described in Haj et al.^32^).

The following plasmids were constructed for this study. Plasmid pJM24 (mTFP1-PTP1Btail) was constructed by fusion PCR of mTFP1 with PTP1Btail (35 C-terminal amino acids of PTP1B) and insertion into a pcDNA backbone. Plasmid pAK63 (mCherry-PTP1Btail) was constructed by swapping mCherry for mTFP1 in pJM24. Plasmids pAK115 (mCherry-PTP1Btail^VCFH^), Plasmids pAK99 (mCherry-PTP1Btail^ΔHALS^), pAK93 (mCherry-PTP1Btail^Scr^), pAK92 (mCherry-PTP1BtailC), and pAK91 (mCherry-PTP1BtailM) were constructed by PCR of mCherry with overlapping 5′ (NheI-containing) and 3′ (EcoRI-containing) primers that included the entire respective coding regions for the amino acid sequences of PTP1Btail^VCFH^, PTP1Btail^ΔHALS^, PTP1Btail^Scr^, PTP1BtailC, and PTP1BtailM (Fig. 6). Plasmid pAK101 (mCherry-PTP1Btail^R428E^) was constructed by double point mutation of the codon for arginine in pAK63 (AGG→GAG). Plasmid pAK110 (mCherry-PTP1Btail^F429R^) by double point mutation of the indicated codon for phenylalanine (TTC→CGC). Plasmid pAK108 (mCherry-PTP1Btail^N412I^) was constructed by point mutation of the indicated codon for asparagine (AAC→ATC). Plasmid pAK47 (mCherry-PTP1B^D/A^-ER) was constructed by replacement of mTagBFP with mCherry-PTP1B^D/A^ in mTagBFP-Sec61β (gift from R. Stricker and E. Zamir). Plasmid pAK49 (mCherry-PTP1B^D/A^-OMM) was constructed by replacement of mTagBFP with mCherry-PTP1B^D/A^ in mTagBFP-CytB5mito (gift from R. Stricker and E. Zamir, contains an OMM-targeting mutant of the tail anchor of CytB5^73^). Plasmid pAK83 (mCherry-PTP1B^D/A^-MAT) was constructed by replacement of mTagBFP with mCherry-PTP1B^D/A^ in COX8a-mTagBFP (gift from R. Stricker and E. Zamir). Plasmid pAK94 (mCherry-PTP1B^D/A^-IMS) was constructed by replacement of N-terminal COX8a sequence in the plasmid mCherry-PTP1B^D/A^-MAT with the N-terminus of the IMS-localizing protein Smac/DIABLO^77^.

### Construction of the Yeast Strains

For preparation of competent yeast and their transformation through homologous recombination of PCR products, we followed standard protocols^88^. For homologous recombination-based insertion into the chromosomal leu2 locus of the wild-type strain ESM356-1^91^ we used appropriately modified versions of the primers ISce1-Nat-A and ISce1-Nat-B (generating the leu2Δ0 deletion) described in Khmlenskii et al.^92^ PCR based insertion of the following plasmids generated the corresponding yeast strains: yemCitrine-PTP1Btail (pAK51, sAK199), yemCitrine-Ysy6tail (pAK72, sAK238), yemCitrine-Sec22tail (pAK86, sAK264), yemCitrine-PTP1BtailM (pAK87, sAK257), yemCitrine-PTP1BtailC (pAK88, sAK258), yemCitrine-PTP1Btail^R428E^ (pAK104, sAK277) and yemCitrine-PTP1Btail^F429R^ (pAK111, sAK281). For C-terminal chromosomal tagging of Ste2, Cox4, Sec7, Cwp2 with mCherry in the indicated strains (sAK201, sAK202, sAK203, sAK204, sAK272, sAK273, sAK274), we used plasmid pFA6a-mCherry-KanMX (gift from M. Knop) and appropriate S3 and S2 primers^88^ for each targeted locus. For deletions of specific insertion pathway proteins in the indicated strains in Table 2, we used appropriately designed S1 and S2 primers^88^ for PCR of the selection factor-containing plasmids pFA6a-klUra3 (gift from M. Knop), pFA6a-HIS3-Mx6 (gift from M. Knop), and pFA6a-hphNT1^88^.

For transformation of the plasmid pAK100 (p415-yemCitrine-PTP1Btail) into the strains RH2881 and RH6829^59^ to respectively generate the strains pAK275 and pAK276, we used the standard lithium acetate-based protocol^93^.

### Transfection, Staining, Fixation, and Immunostaining

Fixation, staining, and immunostaining of the multiple mammalian cell lines shown in Fig. 1 was carried out as follows. COS-7 cells (African green monkey fibroblast-like kidney cells), BJ Fibroblast cells (human foreskin), HeLa cells (human cervical cancer), MCF7 cells (human breast cancer), MDCK cells (Madin-Darby canine kidney), and HepG2 cells (human liver carcinoma) were cultured in growth medium: Dulbecco modified Eagle medium (DMEM; PAN) containing 10% fetal bovine serum (FBS), 1% L-glutamine and 1% non-essential amino acids (NEAA). After staining of the cells with MitoTracker Red CMXRos (250 nM; Invitrogen) for 15 minutes at 37°C, they were washed once with phosphate-buffered saline (PBS) and fixed with 4% paraformaldehyde in PBS (pH 7.5) for 5 minutes at room temperature. The fixed cells were washed three times with tris-buffered saline (TBS), permeabilized with 0.1% Triton X-100 in TBS for 5 minutes at room temperature and then washed again three times with TBS. Blocking was achieved by incubation with 2% bovine serum albumin (BSA) in PBS for 30 minutes at room temperature. Next, the primary antibody PTPase 1B (Ab-1) mouse mAB (Fg61G) (Calbiochem, 1:100 dilution) was applied for 60 minutes at room temperature. Unbound antibody was removed by washing three times with PBS. Secondary antibody incubation was performed for 30 minutes at room temperature with Alexa-Fluor-488 chicken anti-mouse IgG (Invitrogen, 1:200 dilution). Finally, cells were washed three times with PBS followed by their observation with confocal microscopy.

COS-7 cells in Fig. 2 were transiently transfected with a plasmid containing mTagBFP-Sec61 using Fugene6 (Promega) and then fixed and stained as described above. COS-7 cells shown in Figs. 3, 4, 11, 12, 13 and 15 were transfected using Fugene6 (Promega) with the plasmid constructs detailed in the figure captions before their fixation as described above.

COS-7 cells in Figs. 5 and 16–20, as well as Movies 1 and 2, were transiently transfected with the indicated constructs detailed in the figure captions. After 6–8 hours of transfection, the medium was exchanged for growth medium (as described above, Fig. 5 and Movie 1) or starvation medium (DMEM containing phenol red plus 0.1% BSA, Figs. 16–20 and Movie 2) overnight. Immediately before live cell monitoring with fluorescence microscopy (described below), the medium was exchanged for imaging medium (low-bicarbonate DMEM without phenol red; PAN). Stimulation of the starved cells was achieved by addition of EGF at a concentration of 100 ng/mL.

### Confocal Microscopy

All fluorescence images shown in this manuscript (aside from the widefield images in Fig. 5, see section Electron Microscopy below) were obtained using an Olympus Fluoview^TM^ FV1000 confocal microscope (Olympus Life Science Europa, Hamburg, Germany) equipped with an integrated module for time-domain lifetime measurements (PicoQuant GmbH, Berlin, Germany) and a custom-made environmental chamber maintained at 37 °C for the live cell experiments (as the cell media contained HEPES, an additional CO_2_ environment was not necessary). The pinhole was set to 100 μm (roughly one Airy unit) for all images. Continuous 458/488/515 nm lines were produced by an Argon ion laser (Melles Griot, Albuquerque, New Mexico) with an additional 561 nm line was produced by a diode-pumped solid state laser (Melles Griot) in an epifluorescence setting. The integrated module for lifetime measurements consisted of additional five separate pulsed diode lasers (405/440/470/510/532 nm) controlled by a PDL828 “Sepia II” driver. Dichroic elements from Chroma (Rockingham, VT) or Omega Optical (Brattleboro, VT) for detection of the emitted light were chosen with at least 10 nm distance from the relevant excitation wavelength and sufficient distance from the emission profiles of redder fluorophores that may have also been present in the cells. Individual photon arrivals were detected using a SPAD (PDM Series, Micro Photon Devices, Bolzano, Italy) and were recorded by a PicoHarp 300 TCSPC module that could be operated up to a maximum of roughly 10^6^ counts/s. Optimal settings for measuring cells that express all four of the following fluorophores would correspond to the following configuration for our setup: mTagBFP (EX: 405 nm, EM: 420–460 nm), mTurquoise/mTFP1 (EX: 458 nm, EM: 470–490 nm), mCitrine (EX: 510 nm, EM: 524–550 nm), and mCherry (EX: 561 nm, EM: 570–625). All images were obtained at a resolution of either 512 × 512 or 256 × 256 and were often additionally linearly contrasted and cropped (and, where indicated, overlaid) within FIJI^94^ to generate the final displayed images.

### Fluorescence Lifetime Imaging Microscopy

Fluorescence Lifetime Imaging Microscopy (FLIM) images were obtained and analysed as previously described^89^. For the FLIM images shown in Figs. 17–20, pulsed light at 510 nm was used to excite mCitrine in cells expressing ErbB1-mCitrine with single emitted photons between 524–550 nm detected using a SPAD and recorded at up to 10^6^ counts/s (see the section above on Confocal Microscopy). The time-correlated single photon counting (TCSPC) histogram for the entire continuous recording (2 min before EGF stimulation plus 14 minutes after) was used to fit (by χ^2^ minimization) the lifetimes of isolated donors (*τ_1_*) and donors undergoing FRET with the acceptor (*τ*_2_), with single pixel FRET fractions α (along with the constant background) fit using “maximum fidelity”^89^.

### Pulsed Interleaved Excitation

For Movie 1, we employed pulsed interleaved excitation (PIE) to quantitatively assess the colocalization of mTurquoise-PTP1B and mCherry-PTP1Btail across live COS-7 cells. PIE was carried out using our Olympus/PicoQuant FLIM setup (described above). Briefly, pulses of 440 nm and 532 nm light were alternated with a 25 ns pulse interval. The emission light was split using a 560 nm dichroic into two SPADs, one with a 460–500 nm emission filter for collecting the mTurquoise fluorescence (SPAD1) and one with a 570–625 nm emission filter for collecting the mCherry fluorescence (SPAD2). As the mTurquoise fluorescence following the 440 nm excitation pulse will also be present in SPAD2, the events required further filtering, which was performed using a modified version of our custom-written analysis code *p*FLIM^89^. Only events in SPAD1 following the 440 nm pulses were retained (mTurquoise-specific fluorescence). Similarly, only events in SPAD2 following the 532 nm pulses were retained (mCherry-specific fluorescence). This filtering removed the crosstalk between the two channels (as can be seen from our negative control cells in Movie 1, for which the Golgi marker GalNAcT2-mTurquoise was coexpressed with mCherry-PTP1Btail). The final events in each SPAD were then used to generate the individual frames for each fluorophore shown in Movie 1. The red/green overlay images were generated using FIJI^94^ and provide a quantitative and robust measure of the instantaneous ratio of each fluorophore in each pixel.

### Electron Microscopy

Photo-oxidation and DAB precipitation experiments were carried out as previously described^95^. For each photo-oxidation experiment COS7 cells transfected using Fugene6 (Promega) with plasmids containing either mTurquoise-PTP1B or mTFP1-PTP1Btail were briefly washed with prewarmed PBS (phosphate-buffered saline, pH 7.4) and fixed by a mixture of glutaraldehyde (0.5%) and paraformaldehyde (4%) in PBS (Sigma, Germany) for 1 hour. Cells were rinsed 3x with PBS and incubated with the blocking solution containing 100 mM glycine and 100 mM potassium cyanide in PBS for 40 min to reduce nonspecific DAB polymerization. Widefield images for each experiment were obtained with a Zeiss Axio Observer.Z1 inverted microscope (Carl Zeiss, Oberkochen, Germany) fitted with a 63x planapochromat oil objective. For the subsequent photo-oxidation experiments, cells were washed with TBS (TRIS buffered saline, pH 7.4) three times, incubated in freshly oxygenized solution of 2 mg/ml DAB (3,3’-diaminobenzidine tertahydrochloride (Polysciences GmbH, Eppelheim, Germany)) in TBS. The region of interest was imaged with minimal lamp intensity to prevent fluorophore bleaching. Then, the specimen was illuminated with appropriate filter settings (BP excitation filter 436/20 for mTurquoise and mTFP1) using the full power of the 100 W mercury lamp. Photobleaching stopped when a light brownish precipitate was clearly observed with transmission light microscopy at the mitochondria. Then, cells were rinsed with distilled water for 3 min each and subsequently postfixed with 1% osmium tetroxide reduced by 1.5% potassium ferrocyanide for 30 minutes on ice. The specimens were washed with distilled water for 3 min each and dehydrated with graded ethanol series (50%, 70%, 90%, 3x 100%) for 3 minutes each. Finally, cells were embedded in Epon812 (Serva, Eppelheim, Germany) and polymerized at 60°C overnight. The area of interest was trimmed and cut on the Ultracut S microtome (Leica Microsystems, Bensheim, Germany) to get 60-nm slices. The samples were examined on a JEOL JEM-1400 transmission electron microscope (120 kV, JEOL Germany, Eching, Germany) equipped with a 4k CCD camera (TVIPS GmbH, Gauting, Germany).

## Acknowledgements

The excellent technical support of S. Dongard, K. van Eickels, O. Hofnagel, and A. Langerak (MPI of Molecular Physiology) is gratefully acknowledged. ME and MG were supported by the German Ministry of Education and Research (BMBF Grant NanoCombine). We thank M. Knop (Heidelberg U.) and H. Riezman (U. of Geneva) for generous gifts of yeast cells, and M. Knop (Heidelberg U.) and J. Ibach/J. Luig/R. Stricker/P. Verveer/K. van Eickels/E. Zamir (MPI of Molecular Physiology) for generous gifts of plasmids. We thank P.I.H. Bastiaens for his support.

## Competing Interests

All authors declare no competing interests.

---

**Movie 1.** Movie of mTurquoise-PTP1B and mCherry-PTP1Btail. COS-7 cells expressing mTurquoise-PTP1B and mCherry-PTP1Btail (top row) or GalNAcT2-mTurquoise and mCherry-PTP1Btail (bottom row) were confocally imaged using pulsed-interleaved excitation (see Materials and Methods). The cell was continuously tracked for 6 minutes (10 seconds/frame; 60 frames total). The width of the images corresponds to either 132.5 μm (top row) or 151.4 μm (bottom row).

**Movie 2.** Movie of the FRET-based interaction of ErbB1-mCitrine with mCherry-PTP1B^D/A^ as measured with FLIM. Each frame corresponds to 2 minutes. The individual frames before stimulation (-2 min to 0 min) and after stimulation (2 min to 4 min, 12 min to 14 min) are identical to those shown in the fourth row of Fig. 17. Color scale indicates the FRET fraction α (see Fig. 17).

